# Growth-rate dependency of ribosome abundance and translation elongation rate in *Corynebacterium glutamicum* differs from *Escherichia coli*

**DOI:** 10.1101/2021.04.01.438067

**Authors:** Susana Matamouros, Thomas Gensch, Martin Cerff, Christian C. Sachs, Iman Abdollahzadeh, Johnny Hendriks, Lucas Horst, Niklas Tenhaef, Stephan Noack, Michaela Graf, Ralf Takors, Katharina Nöh, Michael Bott

**Affiliations:** Institute of Bio- and Geosciences, IBG-1: Biotechnology, Forschungszentrum Juelich, Juelich, Germany; Institute of Biological Information Processing 1 IBI-1: Molecular and Cellular Physiology, Forschungszentrum Juelich, Juelich, Germany; Institute of Biochemical Engineering, University of Stuttgart, Stuttgart, Germany

## Abstract

The growth rate µ of bacteria depends on the protein synthesis capacity of the cell and thus on the number of active ribosomes and their translation elongation rate. The relationship between these fundamental growth parameters have only been described for a few bacterial species, in particular *Escherichia coli*, but are missing for most bacterial phyla. In this study, we systematically analysed the growth-rate dependency of ribosome abundance and translation elongation rate for *Corynebacterium glutamicum*, a gram-positive model species differing from *E. coli* by a lower growth temperature optimum and a lower µ_max_. Ribosomes were quantified via single-molecule localization microscopy (SMLM) using fluorescently tagged ribosomal proteins and via RNA/protein ratio. Both methods revealed a non-linear relationship with little change in ribosome abundance below µ = 0.4 h^-1^ and a steep increase at higher µ. Unlike *E. coli*, *C. glutamicum* keeps a large pool of active ribosomes at low µ, but the translation elongation rate declines from ∼9 amino acids s^-1^ at µ_max_ to <2 aa s^-1^ at µ < 0.1 h^-1^. A model-based approach shows that depletion of translation precursors at low growth rates can explain the observed decrease in translation elongation rate. Nutrient up-shift experiments support the hypothesis that maintenance of excess ribosomes during poor nutrient conditions enables *C. glutamicum* to quickly restart growth when conditions improve.

## Introduction

*Corynebacterium glutamicum* is a non-pathogenic, rod-shaped, Gram-positive bacterium, used for the production of amino acids in the million-ton-scale, but also for synthesis of various other metabolites and of proteins [1-6]. Due to its outstanding importance in industrial biotechnology and non-pathogenic status, *C. glutamicum* is extensively used as model organism for the development of novel product platforms as well for studying selected topics that are relevant for pathogenic Actinobacteria, such as *Corynebacterium diphtheriae* and *Mycobacterium tuberculosis*. As a result of its widespread use as microbial cell factory, *C. glutamicum*’s metabolism is one of the most studied in bacteria [7, 8]. Fundamental physiological properties such as growth rate can have an important impact on the celĺs metabolism and its productivity in bioprocesses [9, 10]. Therefore, knowledge of the factors that determine and limit the growth rate are of high interest both from a systems biological and a biotechnogical view.

Our understanding of bacterial physiology relies on a few phenomenological growth laws [11, 12] that describe how certain parameters relate to the growth rate. One of these parameters is the cellular ribosome abundance. As growth rate increases, usually in response to better nutrient quality, so does the ribosomal mass fraction. Ribosomes are large multimeric RNA-protein complexes whose synthesis is tightly controlled to avoid loss of fitness due to resource misallocation. A linear correlation between ribosome abundance and growth rate (Rb/µ) has been reported for several organisms, which highlights the importance of protein synthesis to cellular growth [13]. This proportionality relies on ribosomes translating at their maximal speed [14]. Growth rate therefore becomes limited by the number of active ribosomes present in the cell. However, it is known that this correlation is only observed for moderate to fast growth rates (µ > 0.35 h^-1^) [11], as a continuous linear decrease in the number of ribosomes as growth slows down would leave the cells with too few ribosomes to restart growth [12]. Most comprehensive studies on this subject have been performed on the fast-growing model organism *Escherichia coli*. In *E. coli* the translation rate decreases by about 50% from fast to very low growth rates [11, 15-18] denoting that ribosomes are not always working in saturating substrate conditions. The aminoacyl-tRNA pools as well as elongation factors and GTP may become limiting in certain conditions, such as nutrient deprivation or by slow diffusion in the crowded cytoplasm [19-21], leading to a decrease in the elongation rate. Recently, Dai *et al.* elegantly showed that *E. coli* is able to maintain faster-than-expected translation elongation rates during slow growth via significant reduction of the pool of active ribosomes [18].

In this study, we systematically examined the ribosome distribution, abundance and activity for *C. glutamicum* across different growth rates. We found important differences in the fraction of active ribosomes during slow growth between *C. glutamicum* and *E. coli* that may reflect distinct evolutionary strategies for coping with periods of nutrient deprivation. Furthermore, quantitative knowledge of these parameters is important to broaden the current *E. coli*-centred view of these key cellular functions, to enable better growth models, and to identify approaches for improving growth-coupled product formation for this important biotechnological host.

## Results

### Synthesis dynamics and spatial localization of ribosomes in *C. glutamicum* cells

To study cellular ribosome content, distribution and dynamics at different growth rates and growth stages (exponential and stationary), the *C. glutamicum* ATCC 13032 (wt) strain was engineered to chromosomally encode for translational fusions between two ribosomal proteins (bL19 and uS2) and the fluorescent proteins EYFP and PAmCherry. These protein fusions substitute for the native bL19 and uS2 proteins. Synthesis of bL19-EYFP and uS2-PAmCherry was confirmed by western blot analysis of cell extracts (Fig. S1). The resultant dual-labelled fluorescent strain, SM34 (bL19-EYFP, uS2-PAmCherry), grew similarly to the wt strain (Fig. 1a), suggesting that the two fusion proteins were successfully incorporated into the 50S and the 30S subunits, respectively, and did not interfere with ribosome functionality under the selected conditions. Online measurement of EYFP fluorescence in a BioLector cultivation system allowed us to follow ribosome synthesis dynamics during growth, namely that of the large ribosomal subunit. While no fluorescence was detected for the wt strain, for SM34 a fast increase in specific fluorescence (ratio of absolute fluorescence and cell density) was observed upon entry into exponential phase followed by a decrease in mid-exponential phase (Fig. 1a). This early burst in ribosome synthesis is very similar to the one described for *E. coli* [22, 23].

**Figure 1.**
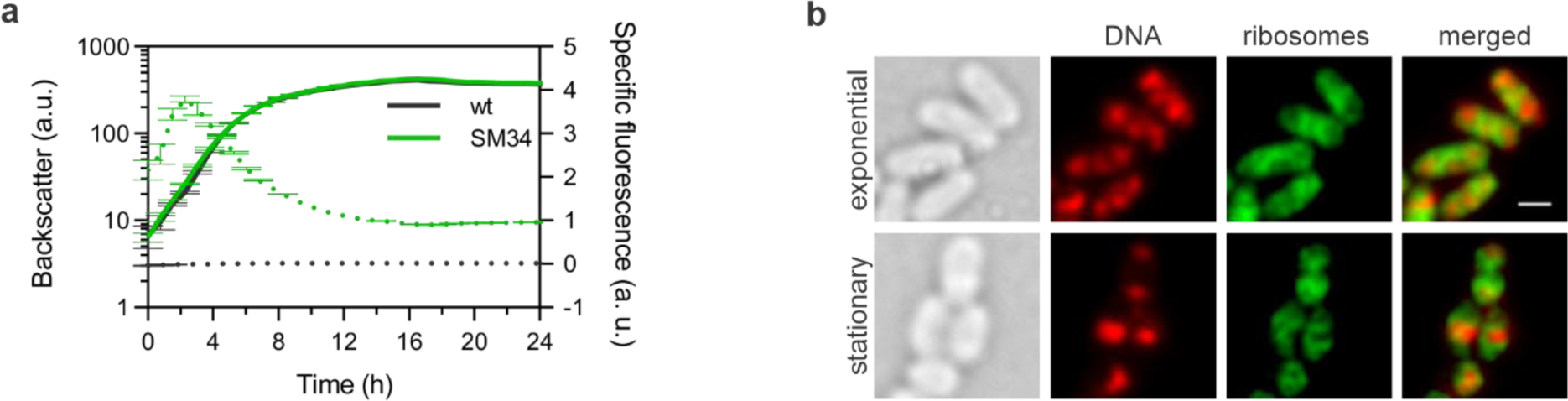
*C. glutamicum* strains carrying fluorescently-tagged ribosomes show wt-like growth behaviour and ribosome-associated fluorescence. a – Growth curve (solid lines) and specific fluorescence (dotted lines) of the wt (grey) and SM34 [(bL19-EYFP uS2-PAmCherry), green] strains from a BioLector cultivation in BHI+GLU medium. Shown is the mean and standard deviation of biological triplicates. **b** – DNA and ribosome distribution of exponential and stationary *C. glutamicum* SM30 (bL19-EYFP) cells cultivated on BHI+GLU. DNA was stained with SYTOX Orange (red). Scale bar = 1 µm.

In some bacterial species [24-29], ribosomes are preferentially found excluded from the highly condensed chromosomal DNA regions, the nucleoids. Therefore, to further verify that the observed fluorescent signal was ribosome-specific, wide-field fluorescence microscopy was performed to examine ribosome and nucleoid localization in exponentially growing and stationary *C. glutamicum* cells. The bL19-EYFP protein was found to be highly abundant in the cell (Fig. 1b), as expected for a ribosomal protein. In addition, although bL19-EYFP was present throughout the cytoplasm it exhibited lower density regions that correlated negatively with DNA-rich regions. In exponentially growing cells, a few DNA-rich regions can be distinguished whereas in stationary phase cells, the DNA is usually found in one highly condensed area (Fig. 1b). In both cases, ribosomes are found surrounding the DNA. This spatial separation suggests that bL19-EYFP is incorporated into ribosomes that due to their large molecular size are less likely to diffuse into the condensed nucleoid areas.

### Ribosome quantification at the single-cell level by single-molecule localization microscopy of *C. glutamicum* cells growing at different growth rates

To quantify ribosome abundance in single cells, a single-molecule localization microscopy (SMLM) method was developed in which uS2-PAmCherry fluorescent molecules were quantified and taken as proxy for the amount of 30S ribosomal particles. For these experiments we chose to work with PAmCherry as it allows for controlled fluorescence activation necessary for quantification [30]. *C. glutamicum* SM34 was grown in different growth media and/or conditions to yield a range of exponential growth rates (µ = 0.20 h^-1^ to µ = 0.49 h^-1^). Immediately after sampling, cells were washed, fixed and mounted on a home-built wide-field fluorescence microscope with single molecule sensitivity [31]. Cells were illuminated with low power 405 nm light (for photoactivation) and high power 561 nm light (for single molecule fluorescence initiation) until all uS2-PAmCherry proteins were excited and detected (see Materials and methods).

Clearly distinct growth phase-dependent ribosome localization distribution was observed in the reconstructed SMLM images (Fig. 2a). In samples from exponential growth phase, ribosomes were found throughout the bacterial cytoplasm with a few observable higher density areas. In stationary phase samples, especially in those where cells were cultivated in glucose-containing medium, ribosomes were preferentially found in high density regions localized either at one or both cell poles (Fig. 2a) or, alternatively, mid-cell. Although the nucleoid localization was not verified in these samples, this result suggests that as observed in the wide-field images, the ribosomes are found in areas probably excluded from the highly condensed nucleoid.

**Figure 2.**
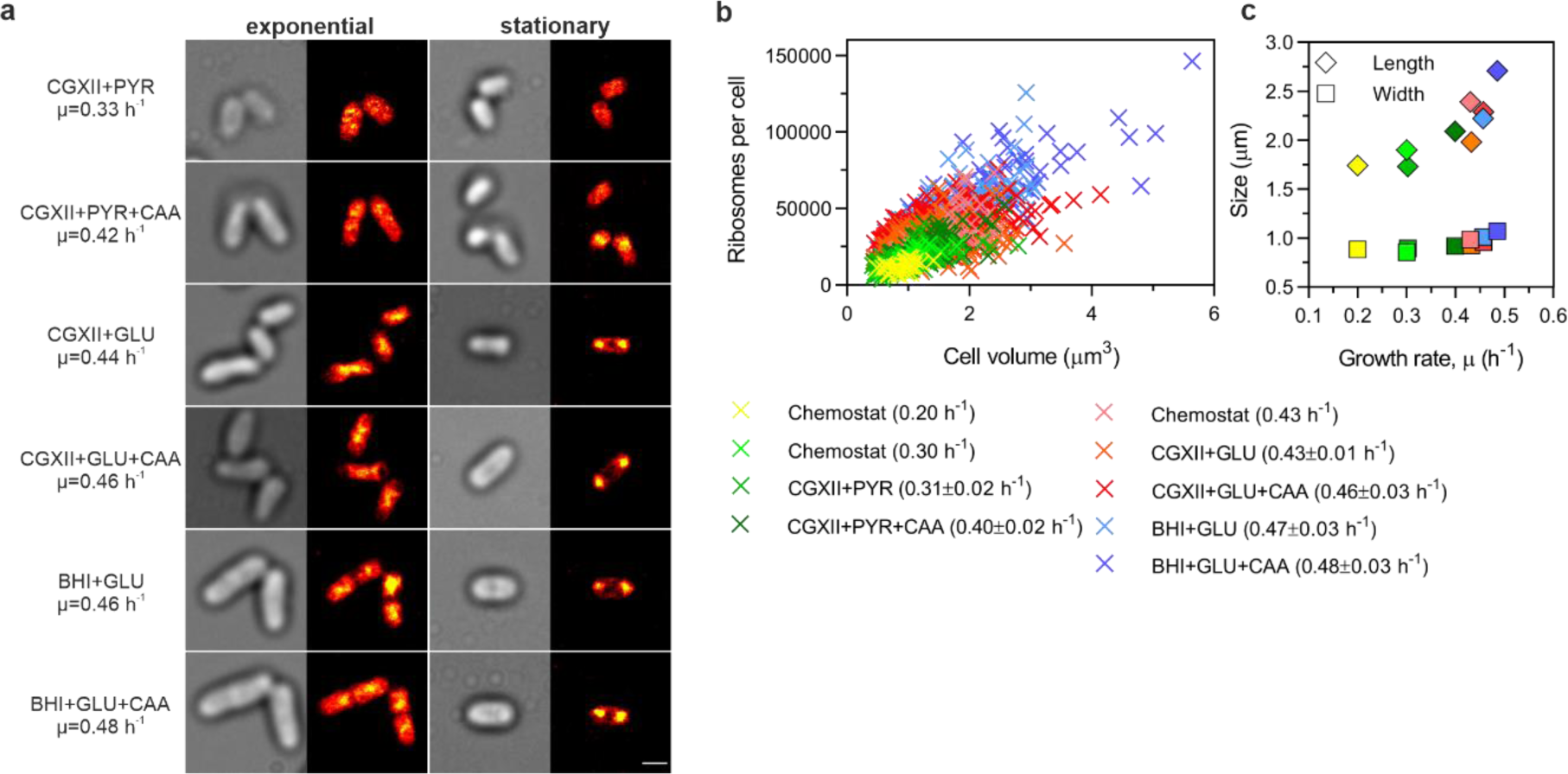
Ribosome number per cell and cell volume increases with growth rate. a – Examples of SMLM images taken from cells cultivated in different media and at different stages of growth. *C. glutamicum* SM34 was cultivated in different media (indicated on the left) enabling increasing exponential growth rates ordered from top to bottom. Samples were taken either in mid-exponential or in stationary growth phase (24 h). Shown are the transmission image (grey) and the corresponding rendered SMLM image (colour) where each dot represents a PAmCherry localization. Higher density regions are shown in yellow whereas darker colours denote lower density regions. Scale bar = 1 µm. **b** – PAmCherry counts (number of ribosomes) determined by SMLM quantification and cell volume are plotted for every individual cell analysed. Cell volume was calculated for every cell following the formula: *V* = π\**W*^2^(*L* − *W*/3)/4 [13]. Each growth condition is colour coded. **c** – The mean cell length (SEM is within 2-7% and approximately the symbol size) and width (SEM is smaller than 2%) is shown for each growth condition (same colour code as in **b**) and plotted against the corresponding growth rate.

To derive per cell ribosome count metrics, emitter molecules previously identified in reconstructed SMLM data via SNSMIL [32] were assigned to individual cells. To this end, the custom software SurEmCo was developed (see Supplementary Note 1). For each condition tested, the total number of fluorescent emitters per cell (uS2-PAmCherry was taken as proxy for ribosome numbers) and the respective cell length and width were determined by the software (Table S1). As already described for other organisms, our results showed that also for *C. glutamicum,* the faster the growth rate the larger the cells are [14, 33-36] and the more ribosomes they contain [24, 37, 38] (Fig. 2b). The observed mean increase in cell volume (from ∼0.9 to ∼2.2 µm^3^) at faster growth rates is mostly due to an increase in cell length from ∼1.7 to ∼2.7 µm (Fig. 2c, Table S1), which results in the so-called *surface area to volume ratio homeostasis* [39, 40]. Also noticeable was the large variation in ribosome numbers and cell volume in cells growing in a certain medium, possibly reflecting the heterogeneity and asynchrony of the culture. In exponentially growing cells, as growth rate increased, the mean ribosome density (ribosomes/µm^3^) increased non-linearly from ∼14,000 at µ = 0.2 h^-1^ to ∼30,000 at µ = 0.48 h^-1^, whereas it remained rather constant in cells in stationary growth phase, irrespective of the medium used for cultivation and the growth rate observed during the exponential phase (Fig. 3a, Table S1). Interestingly, the mean ribosome density in stationary cells was very similar to that of cells growing at slower growth rates (Fig. 3a, Table S1). Control SMLM experiments using a reversely labelled *C. glutamicum* strain, SM55 (uS2-EYFP, bL19-PAmCherry), where in this case bL19-PAmCherry fluorescent molecules were enumerated and taken as proxy for the amount of 50S ribosomal particles, showed very similar ribosome density results for cultivation conditions resulting in similar growth rates (CGXII+GLU and BHI+GLU, Table S1).

**Figure 3.**
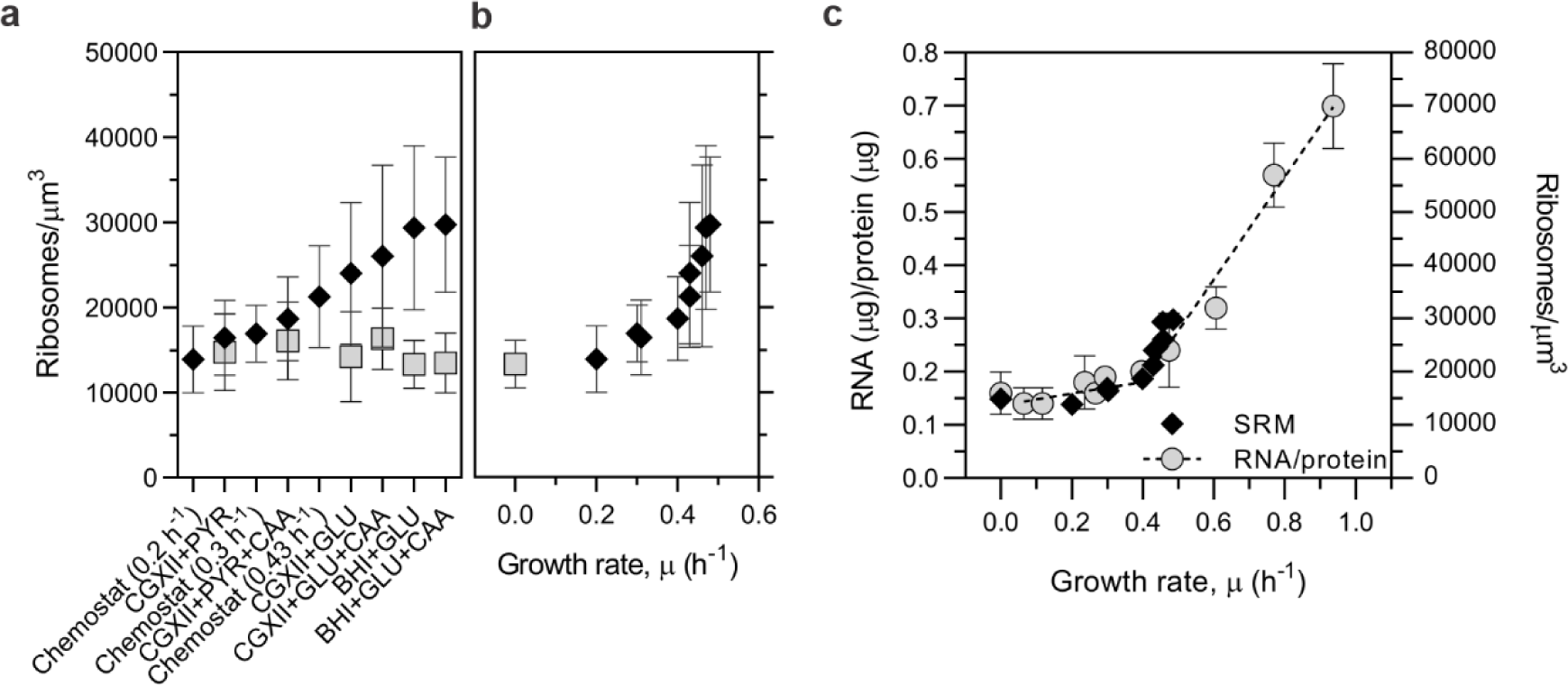
Non-linear Rb/µ correlation in *C. glutamicum*. *C. glutamicum* SM34 was cultivated in different media and conditions yielding different growth rates. Cell samples taken during mid-exponential or in stationary growth phase were imaged by SMLM. **a** – PAmCherry counts (number of ribosomes) per cell volume determined by SMLM quantification for each growth condition tested. Quantification from exponentially growing cultures are represented by black diamonds whereas those from stationary cultures are shown as grey squares. **b** – The values from exponentially growing cultures, shown in **a**, are plotted against the growth rates of the respective cultures (black diamonds). A stationary sample was included as reference for zero growth rate (grey square). **c** – Comparison of the R/P ratio measurements (grey circles) to the SMLM data shown in **b**. Shown are the mean and standard deviation calculated for each condition to which a segmental linear regression was applied. In panel **c**, standard deviation was omitted for the SMLM data set to improve clarity.

### Ribosome quantification at the population level by RNA-protein ratio measurements of. *C. glutamicum* cells growing at different growth rates

The SMLM data revealed a mean ribosome abundance of ⁓15,000/µm^3^ up to a growth rate of 0.2-0.3 h^-1^, which quickly increased to approximately double (⁓30,000/µm^3^) at growth rates of ⁓0.5 h^-1^ (Fig. 3b). To confirm these results and exclude a possible bias in the ribosome quantification method used, we performed classical RNA-protein (R/P) ratio measurements. The R/P ratio is a surrogate and widely used method for the determination of ribosome abundance [13, 37, 41, 42]. To determine the R/P ratio, the *C. glutamicum* wt strain was grown in different media to a yield a range of growth rates and cells were harvested during mid-exponential phase. Unlike *E. coli* or *B. subtilis* that can grow at growth rates >2 h^-1^, *C. glutamicum* wt exhibits a maximal growth rate of ∼0.6-0.67 h^-1^ [43, 44]. In order to analyse cells growing at faster growth rates, an evolved *C. glutamicum* strain (EVO5) [45] was used for some of these experiments (Fig. 3c, Table S2), which reached mean growth rates up to 0.94 h^-1^. The theoretical number of ribosomes per cell can be calculated from the R/P ratio values and the mean colony forming units per OD_600_. Comparison of the ribosome number per cell for *C. glutamicum* wt (see Supplementary Note 2) to that determined by SMLM for strain SM34 grown in the same media (Table S1) showed a good agreement between the two methods and confirmed that the linear Rb/µ correlation is broken for growth rates below 0.4 h^-1^ (Fig. 3c).

### Translation elongation rate and active ribosome fraction

An upward deviation in the R/P ratio at lower growth rates (< 0.7 h^-1^) is also reported for *E. coli* grown at 37 °C [18], but is much weaker than for *C. glutamicum* (Fig. 4a). In *E. coli*, deviation of the R/P ratio from linearity during slow growth is associated with a strong reduction of the active ribosome pool to allow maintenance of high translation elongation rates [18]. To determine if such a process also occurred in *C. glutamicum*, translation elongation rates and derived active ribosome fractions were measured for different growth rates using a fluorescence-based assay (see Materials and Methods and Supplementary Notes 3 and 4).

**Figure 4.**
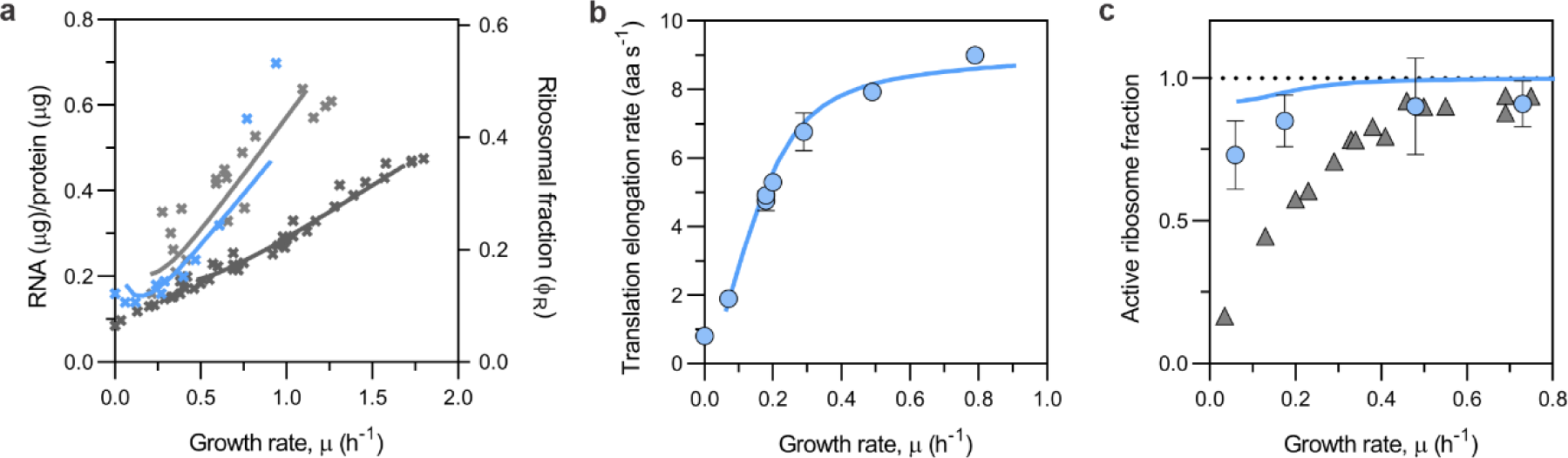
*C. glutamicum* ribosomal fraction, translation elongation rate and inactive ribosome fraction is quite different from that of *E. coli* cultivated at 37 °C. In addition, at low growth rates *C. glutamicum* presents very low translation elongation rates and a high amount of active ribosomes. **a** – R/P ratio or ribosomal fraction of *C. glutamicum* (blue) compared to that of *E. coli* cultivated at either 30 °C (light grey) or 37 °C (dark grey). *E. coli* data was extracted from the literature, for 30 °C [79, 80] and for 37 °C [11, 18, 37, 81] **b** – Translation elongation rate for *C. glutamicum* MB001(DE3) pMKEx2-*eyfp* growing at various growth rates. Depicted are the mean values and standard deviation from at least 2 independent repeats with 3 technical replicates each. **c** – Active ribosome fraction calculated for different growth rates (Supplementary Note 4) for *C. glutamicum* (blue) compared to that of *E. coli* cultivated at 37 °C (grey) extracted from Dai *et al.* [18]. Solid lines in **a**, **b** and **c** were derived from model simulations.

The maximal translation elongation rate determined for *C. glutamicum* (⁓9 amino acids (aa) s^-1^, Fig. 4b) was approximately half that of *E. coli* at 37 °C (17 aa s^-1^ [15, 18]), but very similar to the one determined for *E. coli* at 30 °C (⁓9 aa s^-1^ [46]). The lower translation rates at 30 °C for both organisms are well reflected in the steeper slopes of the linear Rb/µ correlation (Fig. 4a) [13]. *C. glutamicum* grows optimally at 30 °C and therefore that was the temperature at which cells were grown for all assays presented in this work. For *E. coli*, the temperature of choice is usually 37 °C. However, it has been described that also for *E. coli* a higher ribosome abundance is required to support protein synthesis when cells are grown at a lower temperature due to a lower peptide elongation rate, which follows an Arrhenius kinetics for growth temperatures between 23-44 °C [46].

In contrast to *E. coli* grown at 37 °C [18] (no data is available for *E. coli* grown at 30 °C), in *C. glutamicum* the translation elongation rates reached very low values as growth rate decreased (Fig. 4b and S6b). As mentioned above, *E. coli* is able to maintain considerable translation rates during slow growth by reducing the fraction of active ribosomes to under 20% [18] (Fig. S6d). In *C. glutamicum*, however, the fraction of active ribosomes remained above 70% for all conditions tested (Fig. 4c). The higher fraction of active ribosomes probably results in depletion of precursors needed for translation and consequent decrease of the elongation rates observed for *C. glutamicum* towards low growth rates. These results suggest that active ribosome inactivation e.g., via hibernation mechanisms, may not be as relevant for *C. glutamicum* as for *E. coli* under the conditions tested.

### Growth rate and ribosome abundance before and after nutrient upshift

To exclude that the high abundance of ribosomes as well as the marked decrease in the translation elongation rate observed at slow growth rates for *C. glutamicum* was due to the accumulation of defective or misassembled ribosome particles, nutrient upshift experiments were performed. For this purpose the SM34 strain was used, enabling simultaneous measurement of growth (as backscatter) and ribosome production (as fluorescence) in a BioLector cultivation system. Cells were initially grown in a poor carbon and energy source, using either glutamate or ethanol. Once cells reached the exponential growth phase, the preferred carbon source, glucose was added (Fig. 5a and 5b, t=0, vertical dashed line). Growth on glucose and glutamate or glucose and ethanol is known to be diauxic in *C. glutamicum* [47, 48]. Despite different initial growth rates on glutamate and ethanol (0.06 h^-1^ and 0.17 h^-1^, respectively) ribosome accumulation followed very similar kinetics (Fig. 5c). The instantaneous growth rates converged to a maximum of ⁓0.5 h^-1^ 5 h post-transition, which is close to the normal growth rate of *C. glutamicum* when cultivated in CGXII supplemented with glucose as sole carbon source. Immediately after glucose addition there is a very rapid initial jump in the instantaneous growth rate followed by a second slower phase where ribosomes are produced and accumulate to support translation at post-shift rates as evidenced by the progressive increase in specific fluorescence (Fig. 5c) observed ⁓30 min after the upshift. EYFP has a maturation time of 34.0 ± 4.3 min in live *E. coli* cells cultivated at 32 °C [49]. Therefore, ribosome production probably started very soon after glucose addition.

**Figure 5.**
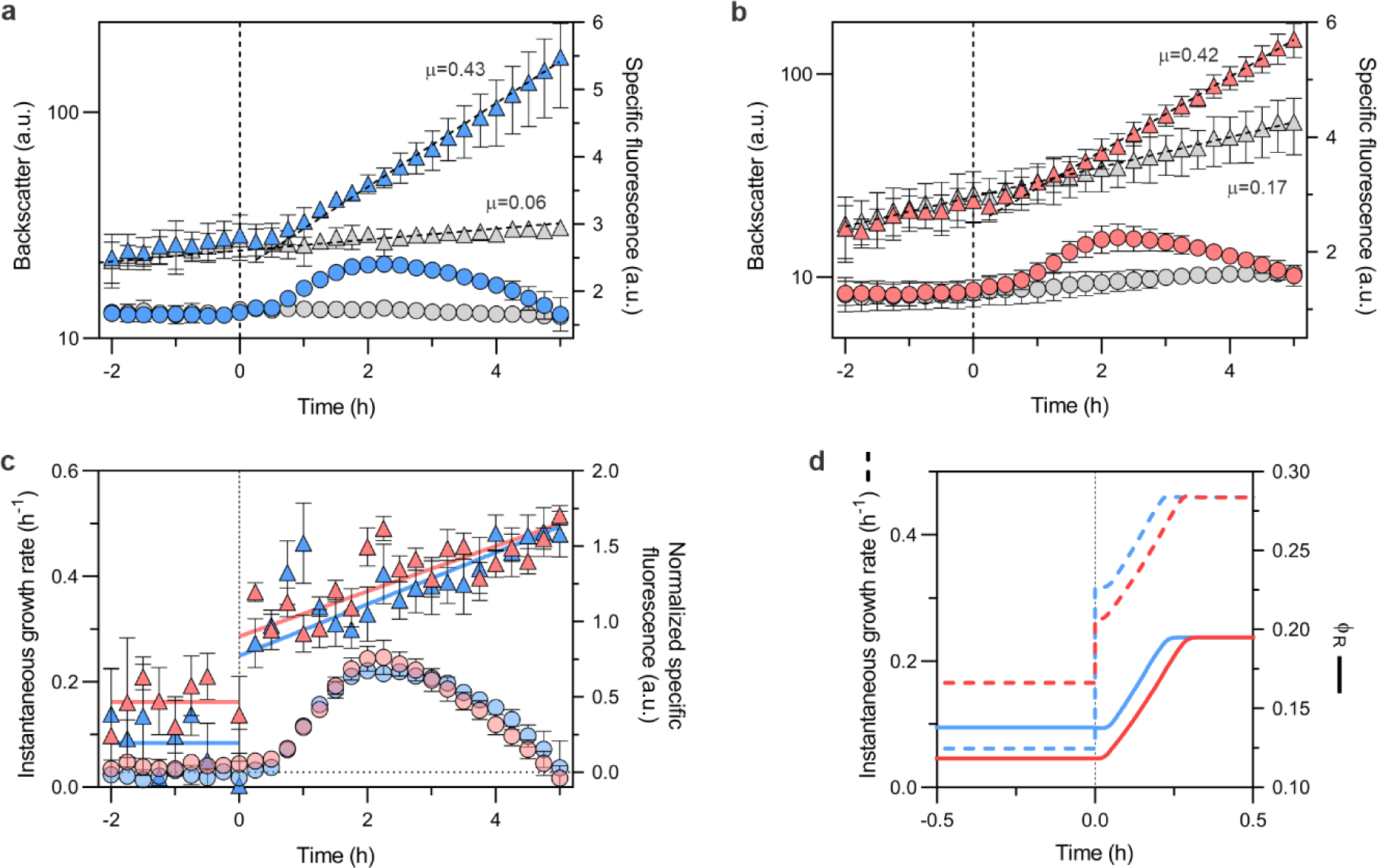
Nutrient upshift results in increased growth rate accompanied by an increase in ribosome abundance. In the upshift experiments cells were initially grown in either CGXII + glutamate (**a**) or CGXII + ethanol (**b**), at t=0 glucose was added (colour-filled triangles) or the same volume of water was added to control wells (grey triangles). Backscatter (triangles) and EYFP fluorescence were followed throughout the entire cultivation on a BioLector. Ribosome abundance is reflected by the determined specific fluorescence for each time point (circles). Growth rates (µ) are indicated above the growth curves and determined by fitting a linear regression to the relevant backscatter values (discontinuous line). **c** – The instantaneous growth rates (triangles) were calculated from the upshift experiments shown in **a** and **b**. The solid lines show the mean instantaneous growth rate for values < 0 h and a linear regression fit for values > 0 h. The normalized specific fluorescence was calculated by subtracting the specific fluorescence of control wells from that of glucose containing wells (circles). Shown are the mean and SEM values from three independent trials with three technical replicates each. **d** – Upshift experiment simulation (see Materials and Methods, Table S9). Shown are time courses of the instantaneous growth rate *µ* (dashed lines) and the ribosomal fraction *ΦR* (solid lines) before and after increasing the nutrient quality parameter (*kn*) from *kn,glutamate* (blue) or *kn,EtOH* (red) to *kn,glucose* (see Table S9).

These results show that in both conditions there is a pre-existing ribosome overcapacity that allows cells to instantly respond to the improved growth conditions. Therefore, we conclude that the high ribosome abundance observed for *C. glutamicum* during slow growth is not due to the accumulation of non-functional ribosomal particles.

### Mechanistic insights into the Rb/µ correlation in *C. glutamicum*

To reveal deeper insights into the dependency between growth rate and ribosomal fraction and to understand if depletion of translation precursors could result in the very low translation rates observed during slow growth, the *E. coli* coarse-grained self-replicator model by Bosdriesz *et al.* [50] was re-calibrated for *C. glutamicum* (see Materials and Methods). The model formulates the resource-balancing act of cellular investments between ribosome production, translation elongation and growth. It uses translation inefficiency, i.e. the amount of uncharged tRNAs bound to the ribosome, as signal for the regulation of ribosome synthesis via a ppGpp-based regulatory mechanism. As in *E. coli*, in *C. glutamicum* total RNA and growth rate are inversely correlated to the cellular ppGpp concentration [51]. Equipped with this simple regulatory mechanism, the calibrated model was able to accurately describe the nonlinear Rb/µ correlation for *C. glutamicum* over the full growth range (*µ* = 0.07-0.94 h^-1^, Fig. 4a and Supplementary Note 6 for comparison with *E. coli*). From the model, we inferred key rates of translation elongation and ribosome production, i.e. the maximum translation elongation (*kR,max* = 9.3 ± 0.8 aa ribosome^-1^ s^-1^) and *rrn* operon transcription initiation (*vrrn* = 0.3 ± 0.1 -11.1 ± 3.8 *rrn* initiations s^-1^ cell^-1^) rates. Inferred *kR* were in excellent agreement with the experimentally determined values (Fig. 4b), whereas *vrrn* rates were in the same order of magnitude as those calculated from cellular ribosome numbers (Fig. S6c). Furthermore, from the modelling results, a high fraction (91.7-99.8%) of active ribosomes was predicted for the entire range of growth rates (Fig. 4c). In the model this is explained by the fact that the estimated concentration of unloaded tRNAs (Fig. S7b) and thus inactive ribosomes (Fig. S7d) as well as ppGpp (Fig. S7e) only steeply rises at very low growth rates. This results not only in the maintenance of a high fraction of active ribosomes, but also in the associated decrease of translation elongation rates due to depletion of translation precursors such as aminoacylated-tRNAs at slow growth (Fig. S7a), analogous to what was experimentally determined (Fig. 4b and 4c). Therefore the coarse-grained model with its regulatory mechanism for control of ribosome synthesis was able to well reproduce the experimental observations for *C. glutamicum*. Noteworthy, this sets a counterpoint to *E. coli* (37 °C), where it cannot explain the relatively high translation rates during slow growth (*µ* < 0.5 h^-1^, see Supplementary Note 6).

To further analyse the dynamic response to nutrient upshifts we modelled the adaptation of *C. glutamicum* when transferred into more nutritious media (Fig. 5d). In the simulations the progressive increase in ribosome fractions observed in the experimental results (Fig. 5c) were well captured (Fig. 5d) as result of the expected rise in ribosome production rate (*vrrn*, Fig. S8a), which initially overshoots before leveling off at a new steady-state value. Furthermore, transition into a richer medium resulted in the increase in ribosome activity (Fig. S8b), which, due to the presence of the additional translation precursors, caused a jump in translation rate (*kR*) (Fig. S8a) and a consequent step increase in growth rate (Fig. 5c and 5d), probably modulated by the intracellular ppGpp-concentration (Fig. S8c). Finally, to investigate the maximum possible growth rate of *C. glutamicum*, e.g. after a nutrient upshift to a “super-rich” medium, extrapolation of the nutrient quality *kn* (see Supplementary Note 5D) predicted a global maximum specific growth rate ^μ̂^_*max*_ = 0.94 h^-1^, which is very close to the experimentally determined mean growth rate of the laboratory-evolved EVO5 strain (Table S2, [45]).

## Discussion

In this study, we examined fundamental physiological properties, namely the cellular ribosomal localization, ribosome abundance and the translation elongation rate of *C. glutamicum* over a range of growth rates. We show that in *C. glutamicum* there is spatial separation between ribosomes and chromosomal DNA during exponential and stationary growth phases. Compartmentalization of translation can have important physiological implications. The tight organization of chromosomal DNA into spatially separated nucleoids within the cell generates DNA-free areas where the mobility of large complexes such as ribosomes is found to be increased when compared to that of bacterial species where no separation between DNA and ribosomes is observed [29]. Furthermore, it can influence mRNA localization, either close to the gene locus from which the mRNA was transcribed from [52] due to decreased local diffusion or close to the cellular location of the encoded protein(s) [53]. We speculate that the latter may also occur in *C. glutamicum* since the ribosomes are preferentially found in nucleoid-free areas.

To quantify ribosomes in single cells, we developed a SMLM method together with the emitter counter software SurEmCo. This method and analysis may easily be applied to the study of other proteins in this or other organisms. From our estimates, a resting *C. glutamicum* cell in stationary phase possesses ⁓15,000 ribosomes/µm^3^. At the highest growth rate of ⁓0.5 h^-1^ analysed by SMLM, this number doubles to ⁓30,000 ribosomes/µm^3^, which is close to the reported 27,000 ribosomes/µm^3^ for *E. coli* at 30 °C and µ ≈ 0.7 h^-1^ determined by a different super-resolution microscopy method [24]. However, in our SMLM analysis we remarked a steep increase in ribosome abundance above µ ≈ 0.4 h^-1^ and these results were confirmed by R/P ratio measurements of the wt and an evolved faster-growing *C. glutamicum* strain. Such marked deviation of R/P ratio from linearity during slow growth has also been reported for *Aerobacter aerogenes* [42]. In *E. coli*, although present, this deviation is considerably more moderate [18]. However, due to the lack of systematic examination of these parameters during slow growth for other organisms and their implications on ribosome activity, our results are further discussed in the light of what is known for *E. coli* at 37 °C.

As for *E. coli* [18], the inflection point at which the R/P ratio starts to linearly rise with the growth rate is approximately the point at which the translation elongation rate is close to its maximum value. At µ < 0.4 h^-1^, faster growth of *C. glutamicum* is thus mainly achieved by the increase in the rate of protein synthesis rather than by an increase in ribosome abundance. Unlike *E. coli*, for which the translation elongation rate varies only by approximately 2-fold between slow and maximal growth rates [17, 18], for *C. glutamicum* we observed an almost five-fold increase in the translation elongation rate across all steady-state growth conditions at which it was measured (1.9 – 9.0 aa s^-1^). The increase was even more than ten-fold if one considers the translation elongation rate measured in stationary phase (⁓0.8 aa s^-1^). Dai *et al.* show that the translation elongation rate in *E. coli* is kept high due to substantial ribosome inactivation during slow growth [18]. However, for *C. glutamicum* we observe that the fraction of active ribosomes is instead kept high, above 70% for all tested conditions. The marked drop in translation elongation rate could be well simulated by a coarse-grained self-replicator model that uses the amount of uncharged tRNAs bound to the ribosome as signal for the regulation of ribosome synthesis. Ribosome hibernation mechanisms, in which bacteria inactivate a fraction of their own ribosomes to reduce the cost of protein production while waiting for more favourable conditions, have been elucidated for a few organisms [54]. For *Corynebacterium* spp. not much is known on how ribosome activity is modulated nor what the consequences for its physiology are. However, the presence of a large fraction of active ribosomes argues against the relevance of such mechanisms under the conditions examined in this study. A summary of the major results as well as a model of how the ribosome abundance, the translation elongation rate and the active ribosome fraction vary in function of the growth rate in *C. glutamicum* can be found in Fig. 6.

**Figure 6.**
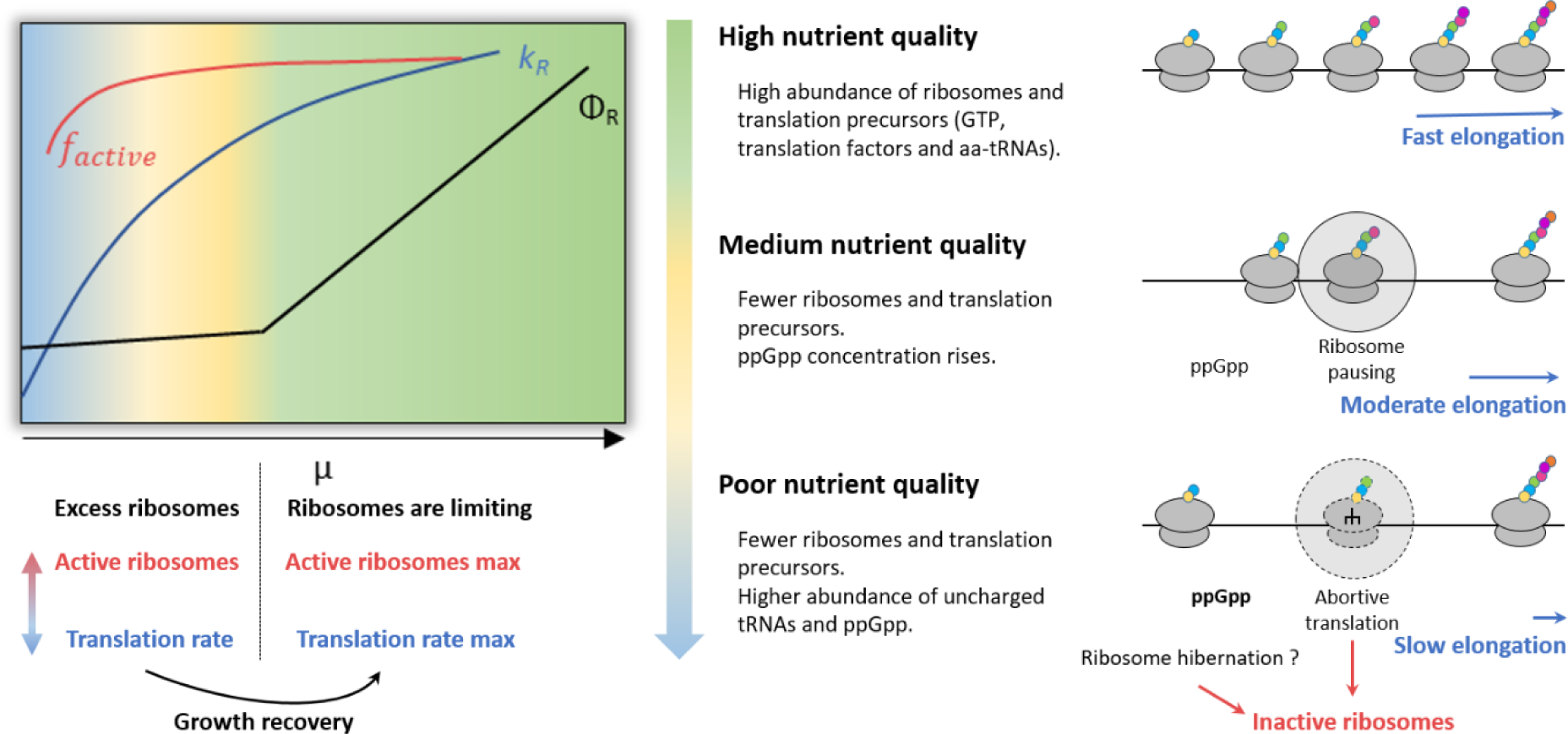
Model of how the ribosome abundance, the translation elongation rate and the active ribosome fraction vary as function of the growth rate in *C. glutamicum*. Under good nutrient conditions there is no shortage of translation precursors. The majority of ribosomes are active and translation elongation reaches close to maximum rates, thus faster growth is achieved by increasing the ribosome fraction of the proteome. This gives rise to the well-established linear Rb/µ correlation [13]. In less optimal conditions, translation precursors become limiting triggering ppGpp synthesis. In some bacteria, this leads to the downregulation of ribosome production [56] as well inhibition of several GTP-dependent enzymes involved in translation such as the initiation factor 2 (IF2) [82] and ribosome biogenesis factors [83]. The decrease in ribosome abundance, translation initiation and ribosome assembly can all lead to a lower elongation rate. In addition, lack or low abundance of aminoacyl-tRNAs can cause ribosome pausing that if not resolved results in abortive translation events [84, 85]. These events lead to premature translation termination of stalled ribosomes and subsequent degradation of the partially translated peptide(s) [85]. Ribosomes involved in abortive translation events are then considered inactive since they do not produce the intended protein. Alternatively, ribosomes can also be kept in an inactive form through ribosome hibernation mechanisms [54]. Significant ribosome inactivation in *E. coli* frees substrates needed for translation and allows substantial translation rates during slow growth [18]. However, under the conditions examined in this study, we found that ribosome hibernation probably does not play an important role in *C. glutamicum*. Instead, we show via a model-based approach that the depletion of translation precursors such as aminoacylated-tRNAs during slow growth is able to reproduce the experimentally determined very low elongation rates measured. Towards lower growth rates, when nutrients are limiting, there is then a balance between the fraction of active ribosomes and their translation elongation rate. Under this regime ribosomes exist in excess to allow for quick growth recovery once conditions improve. How different organisms manage this balance appears to be different and may depend on the environmental niches in which they evolved.

Ribosome biogenesis is tightly controlled in a growth rate-dependent manner [55]. The alarmone (p)ppGpp is a major modulator of the bacterial growth rate, as it regulates ribosome production in response to the nutritional state of the cell [56]. In *C. glutamicum* [51], as in other bacteria [56], the cellular ppGpp concentration is inversely correlated with the growth rate. In addition, most translation-associated genes are transcriptionally downregulated in a (p)ppGpp-dependent manner [57]. However, in contrast to other bacteria such as *E. coli*, where accumulation of uncharged tRNAs strongly activates ppGpp synthesis, in *C. glutamicum* and in mycobacteria strong ppGpp upregulation is only observed upon complete depletion of all carbon-, nitrogen- and phosphorus-containing nutrients [57-59]. Differences in the stringent response triggers have been suggested to reflect adaptations to the distinct lifestyle of each species [60].

Excess ribosomes at slow growth is also described for other bacteria and can be seen as an evolutionary adaptation [61-64]. *C. glutamicum* has been shown to possess at least two chromosome copies across different growth conditions with accumulation of chromosome equivalents in stationary phase [65, 66]. Moreover, in certain conditions granular polyphosphate, an important energy and phosphate reservoir for metabolism and growth, can accumulate to high levels in this organism [67]. These characteristics as well as the presence of a high abundance of ribosomes during slow growth may be seen as adaptations to long periods of starvation caused e.g., by dilute environments as those probably experienced by *C. glutamicum* during evolution as a non-motile soil-dwelling organism. Maintenance of a high ribosome abundance may allow for faster recovery from nutrient deprivation periods. During evolution organisms may have geared either towards growth rate maximization or the ability to survive long periods of starvation [68]. As shown by Mori *et al.*, upon a nutrient upshift strains with large translational overcapacity initially perform well due to the rapid increase in growth rate, however they are ultimately outperformed by the other strains with higher final growth rate [68]. In fact, lag-phase as described for other bacteria such as *E. coli*, e.g. when inoculating fresh medium with an overnight culture cultivated in the same medium composition, are not observed for *C. glutamicum*. In order to induce long lag phases, cells have to be starved for several days [9]. This is suggestive of a large translational overcapacity of *C. glutamicum* during slow growth as observed in the upshift experiments. Cells growing at a growth rate of 0.06 h^-1^ (glutamate) or 0.17 h^-1^ (ethanol) were able to immediately shift to an instantaneous growth rate of ∼0.27 h^-1^ or ∼0.37 h^-1^, respectively, without an apparent transition phase. Since the fraction of active ribosomes for *C. glutamicum* remained fairly high for all conditions tested, the increase in growth rate likely comes from a faster translation elongation rate due to the presence of readily accessible precursors made available by the newly added carbon-source to the medium. In a previous study, we have also shown that in *C. glutamicum* central metabolic enzymes are more abundant than required to support a certain metabolic flux [69]. This ensures a fast supply of amino acid precursors to quickly respond to environmental changes.

Mori *et al.* postulated a fundamental trade-off between growth rate optimization and rapid growth recovery [68]. For *E. coli* it is well established that the former is employed as evolutionary strategy result of the famine-and-feast cycles encountered by the gut microbiota. Here, we provide evidence that the converse is true for *C. glutamicum*. Maintenance of an excess of slow-working ribosomes under poor nutrient conditions allows for a large translational overcapacity that enables quick growth recovery when conditions improve at the cost of a lower maximal growth rate (Supplementary Note 6 and Table S8).

## Materials and Methods

### Bacterial strains and plasmids

The strains used in this study were the *C. glutamicum* wild-type (wt) strain (ATCC 13032) or its derivatives MB001(DE3), a prophage-free strain which allows for IPTG-inducible T7-RNA polymerase-dependent gene expression [70] and EVO5, a laboratory evolved faster-growing strain [45]. *E. coli* DH5α was used in all cloning procedures.

For fluorescent co-imaging of ribosomes and nucleoids, *C. glutamicum* wt was modified via homologous recombination to carry a translational fusion between the ribosomal protein bL19 (RplS) and the EYFP fluorescent protein, giving rise to strain SM30. The *eyfp* gene was amplified from the pEKEx2-*eyfp* [71]. It originates from the pEYFP-C1 (Clontech) and encodes an EYFP protein sequence with two amino acids changes (D130N and D174G) that do not affect its photophysical properties (personal communication Dr. Julia Frunzke). The *rplS* (cg2235) gene along with 1 kb of its upstream region, the EYFP coding gene and 1 kb of the *rplS* immediate downstream region were PCR-amplified. The resulting DNA fragments were subsequently cloned in the above indicated order (1kb_upstream-*rplS*-linker-*eyfp*- 1kb_downstream) via Gibson assembly into the EcoRI-digested pK19*mobsacB C. glutamicum* suicide vector [72]. During cloning the linker sequence 5’-GGCGCTGCTGCTGCTGGC-3’ corresponding to the amino acid sequence GAAAAG was inserted between *rplS* and *eyfp* gene sequences. The resulting plasmid was introduced into *C. glutamicum* wt via electroporation.

Double crossover mutants were identified via single clone selection on kanamycin-containing plates followed by counterselection on sucrose-containing plates as described in detail in [73]. Finally, correct integration was confirmed by colony PCR and Sanger sequencing of the region of interest. For SMLM, strain SM30 was modified via homologous recombination to carry an additional translational fusion, in this case between the ribosomal protein uS2 (RpsB) and the photoactivatable mCherry fluorescent protein (PAmCherry) [74], yielding strain SM34. To construct the uS2-PAmCherry translational fusion the *rpsB* (cg2222) gene along with 1 kb of its upstream region, the PAmCherry encoding gene and 1 kb of the *rpsB* immediate downstream region were amplified by PCR. As above, the resulting DNA fragments were subsequently cloned in the presented order (1kb_upstream-*rpsB*-linker-*PAmCherry*- 1kb_downstream) via Gibson assembly into the EcoRI-digested pK19*mobsacB* suicide vector [72]. During cloning the linker sequence 5’-CAGGAAAGGCGACAGGAG-3’ corresponding to the amino acid sequence QERRQE [24] was inserted between *rpsB* and PAmCherry gene sequences. Double crossover selection and correct integration verification was performed as described above for SM30. To construct strain SM55, which carries the translational fusions uS2-EYFP and bL19-PAmCherry, a similar approach as described above for SM34 was followed. In this case, we started by creating a strain containing the bL19-PAmCherry translational fusion (linker sequence - 5’-GGCGCTGCTGCTGCTGGC-3’ encoding the amino acid sequence GAAAAG) into which the uS2-EYFP (linker sequence - 5’-CAGGAAAGGCGACAGGAG-3’ encoding the amino sequence QERRQE) was subsequently introduced.

To measure the translation elongation rate, the *C. glutamicum* MB001(DE3) strain carrying the reporter plasmid pMKEx2-*eyfp* was used [70].

### Growth medium and cultivation conditions

*C. glutamicum* strains were cultivated either in brain heart infusion (BHI) medium (Bacto^TM^, BD, Heidelberg, Germany) or in CGXII mineral medium containing per liter of distilled water: 1 g K2HPO4, 1 g KH2PO4, 10 g (NH4)2SO4, 5 g urea, 21 g 3-(*N*-morpholino)propanesulfonic acid (MOPS), 0.25 g MgSO4.7H2O, 10 mg CaCl2, 0.2 mg biotin, 10 mg FeSO4, 10 mg MnSO4, 1 mg ZnSO4, 0.3 mg CuSO4, 0.02 mg NiCl2, supplemented with 30 mg of 3,4-dihydroxybenzoate (iron chelator) and a sole carbon source (below). The pH was adjusted to 7.0 with KOH. The different carbon sources were provided at the following final concentrations: 2% (w/v) glucose (GLU), 2% (w/v) sodium acetate (ACE), 2% (w/v) sodium pyruvate (PYR), 2% (w/v) sodium DL-lactate (LAC), 1.1% (v/v) ethanol (EtOH), and 0.9% (w/v) sodium glutamate (GLUT). In a few conditions 0.2% (w/v) casamino acids (CAA) or yeast extract (YE) was added. Whenever necessary, the medium was supplemented with kanamycin (25 µg/ml).

*C. glutamicum* cultures were always cultivated at 30 °C and followed three steps: seed culture, preculture and experimental culture. In the seed culture, single clones were grown overnight (ON) in BHI+GLU and then diluted 1:100 for the precultures, which were cultivated for ⁓18 h in the same medium as the experimental cultures. Finally, the experimental cultures were inoculated at a normalized starting optical density at 600 nm (OD_600_) of 0.5-1.0.

For microscopy, cells were grown in 10 ml of culture medium in 100 ml baffled flasks shaken at 130 rpm. For RNA/protein measurements, cells were grown in 50 ml of culture medium in 500 ml baffled flasks shaken at 130 rpm or in a BioLector cultivation system (m2p-labs, Baesweiler, Germany). For BioLector microcultivations, cells were grown in 800 µl culture volume in a 48-well flower plate shaken at 1200 rpm.

To control for environmental variations such as the supplied medium as well as those inherent to shake flask cultivations, such as oxygen availability and/or pH shifts, samples were also taken from continuous chemostat cultivations. For the chemostat cultivations, a 3 L steel bioreactor (KLF 2000, Bioengineering, Switzerland) equipped with process control (Lab view 2010, National Labs) was used. Continuous dilution was regulated with an advanced controller scheme [45] and three dilution rates were installed determining the exponential growth rate of the culture (µ = 0.2, 0.3 and 0.43 h^-1^). Cells were cultivated in 1.2 L CGXII culture medium supplemented with 2% (w/v) glucose and 0.2% (w/v) CAA at 30 °C and 1.5 bar. pH was controlled at 7.4 via automatic addition of 25% (v/v) NH4OH solution and dissolved oxygen was kept above 70% by setting constant stirrer speeds of 700 rpm and aeration rates of 0.85 L min^-1^. Samples from cells growing at µ = 0.2, µ = 0.3 and µ = 0.43 h^-1^ were immediately prepared for single molecule localization microscopy (see below).

### Wide-field and single molecule localization microscopy

*C. glutamicum* SM30 was cultivated in BHI+GLU until mid-exponential phase (OD_600_ ⁓ 3.0-5.0) or until stationary phase (24 hours). At these time points, 500 nM SYTOX Orange (ThermoFisher Scientific, Waltham, MA, USA) was added and the cultures were further incubated in the same cultivation conditions for 10 min. Samples of 150 µl were collected by centrifugation at 5000 x *g*, washed once in 1 ml 1X phosphate-buffered saline (PBS) and fixed with a 4% (w/v) paraformaldehyde solution in 1X PBS. After 20 min incubation at room temperature the reaction was stopped by centrifuging and resuspending the cells in 1X PBS containing 10 mM glycine. Cells were imaged on 1% (w/v) agarose pads containing 1X PBS mounted on a glass slide.

All fluorescence microscopy measurements were performed with a self-constructed wide-field and TIRF fluorescence microscope with single-molecule sensitivity based on an Olympus IX-71 inverted microscope body. It uses an AOTF (AOTF nC-VIS-TN 1001; AA Opto-Electronic, Orsay, France) to control the throughput of the two continuous wave laser sources that we used, i.e. an Argon-ion laser (514 nm; Coherent Innova 70C, Coherent Inc., Santa Clara, US) for excitation of EYFP and a 561 nm solid state OPSL laser (Sapphire 561-200 CDRH-CP; Coherent Inc.) for excitation of photoactivated PAmCherry or SYTOX Orange. An Olympus ApoN 60x oil TIRF objective (NA 1.49) was used. Excitation and emission light were separated via a multiband dichroic mirror (Di01 R442/514/561; Semrock IDEX Health and Science LLC, Rochester, NY, USA) in combination with a multiple bandpass filter (FF01-485/537/627; Semrock) and an additional single bandpass filter either FF01-609/57 (Semrock) for imaging SYTOX Orange and photoactivated PAmCherry or FF03-525/50 (Semrock) for EYFP imaging. Images were recorded with an EMCCD camera (Andor iXon DU897E-C00-#BV, Oxford instruments, Abingdon, UK) cooled to -75 °C using a resolution of 512 x 512 pixels. The image from the microscope is additionally magnified via an achromatic lens (focal point, 50 mm; AC254-050-A-ML; Thorlabs, Bergkirchen, Germany). By adjusting (motorized) the lens and camera position, the pixel size can be adjusted between 65 and 130 nm/pixel. We used 80 nm pixel size. The camera acquisition time was set to 50 ms in all experiments. It is followed by a 35 ms readout time during which the camera cannot detect photons, i.e., in an image sequence, every image represents 85 ms, with an effective detection time of 50 ms. Wide-field images were calculated as the mean image of a short image series (typically 50) using 1 or 2% of the excitation laser output power.

For single molecule localization microscopy (SMLM), cell samples from mid-exponential or stationary cultures were centrifuged at 5000 x g, washed once in 1 ml 1X phosphate-buffered saline (PBS) and fixed with 4% (w/v) paraformaldehyde as detailed above. A volume of 200 μl of appropriate cell dilutions in 1X PBS was applied to a well of an 8-well chambered micro-slide (µ-Slide 8 Well Glass Bottom (Cat.No: 80827), ibidi GmbH, Gräfelfing, Germany) previously coated with 0.1% (w/v) poly-L-lysine. After 30 minutes incubation at room temperature the micro-slide chambers were washed twice with 1X PBS to remove any unattached cells. The slides were kept at 4 °C for a maximum of 7 days before use. SMLM was performed with the photoactivatable fluorescent protein PAmCherry [74]. PAmCherry cannot be excited at the imaging wavelength of 561 nm, since it absorbs in the near UV/blue spectral region (absorption maximum), fluorescing very dimly in the green spectral region. Photoactivation occurs when applying 405 nm light, upon which PAmCherry is transformed in a complex photochemical reaction [75] to a bright emitting fluorophore with an excitation maximum at 564 nm and an emission maximum at 595 nm. Super-resolved images were obtained by recording a long sequence of wide-field images (50 ms acquisition) with high power (75%) 561 nm laser light under continuous 405 nm diode laser exposure (Cube 405 - 100C; Coherent). As we intended to count all ribosomal PAmCherry fusion proteins present in the cells, we had to ensure that only a few photoactivated PAmCherry molecules per cell were present at a time and were detected as single molecules by excitation at 561 nm. For this reason, we used at the beginning of the SMLM acquisition a very low power of the photoactivating 405 nm light and manually increased the 405 nm laser power to keep a constant, low level of emitters per cell to avoid too many emitting PAmCherry molecules in single images. In fact, in some experiments there was a minute fraction of PAmCherry molecules that were already absorbing at 561 nm before the photoactivating light was applied. Therefore, we first acquired 100-300 single molecule images without any 405 nm light before we started the photoactivation.

Typically, we had 0 to at most 5 PAmCherry molecules per cell emitting at the same time, i.e., in the same wide-field image. In this way we ensured that we do not fail to detect any emitter due to problems of insufficient spatial separation of single molecules in single images as it would occur using SNSMIL or any other standard single molecule localization algorithms [76]. If emitters are too dense and their diffraction-limited representations (circles with diameter of ca. 250 nm) would overlap, SNSMIL would interpret the resulting fluorescence pattern as one single emitter (instead of two or three) and we would have systematically determined too low ribosome numbers.

As mentioned above, we gradually increased the 405 nm light power when recording the single molecule wide-field fluorescence image sequence, once the number of emitters in the current image was rather low (0 or 1 emitter per cell). We continued in this way until a further increase of the 405 nm light power did not generate new emitting single molecules. The recorded image sequences contained 5,000 – 60,000 images and were analysed by SNSMIL, which determined the number, intensities and positions of all single molecule emitters in all cells and in all images of the image sequence. These data were further analysed by an in-house developed single molecule tracking program to correct for two additional problems in the determination of ribosome number per cell (Supplementary Note 1).

### Ribosome counting by SMLM

To derive per cell ribosome count metrics, previously reconstructed snapshot SMLM data was tracked over time, and emitters assigned to individual cells. To this end, a custom software SurEmCo (Super-resolution Emitter Counter) was developed (https://github.com/modsim/suremco). The Python software identified individual cells using a local thresholding method [77] applied to an acquired transmission image. Identified cell areas were used to select all raw emitter positions inside cells as generated by SNSMIL [32]. Emitters (i.e. ribosomes) were tracked throughout the image sequence per cell region, since they might slightly drift over time. Every emitter track with at most one dark frame in between was counted as one emitter to counteract potential biases in emitter counts caused by the presence of the same fluorophore in consecutive frames and the blinking probability of the fluorophore (Supplementary Note 1). Results were visualized in 3D, and metrics provided in a tabulated format. All processed outcomes were manually examined to exclude contributions originating from falsely identified cellular regions or superposed cells.

### RNA extraction

For each condition, the volume equivalent to an OD_600_ of 3-5 per ml of a mid-exponentially growing culture was pelleted in screw cap centrifuge tubes, flash frozen in liquid nitrogen and stored at -80 °C. Pellets were resuspended in 500 µl QIAzol lysis reagent (Qiagen, Hilden, Germany) and incubated at room temperature for 1 min. Cells were disrupted using glass beads (3 x 30 s at 6500 rpm) in a Precellys 24 homogenizer (PEQLAB Biotechnologie GmbH, Erlangen, Germany) with a 5 min incubation on ice in between each disruption cycle. RNA was extracted once with chloroform using phase lock gel tubes (5 PRIME GmbH, Hilden, Germany), precipitated with isopropanol and washed with 70% (v/v) ethanol. RNA pellets were allowed to dry at room temperature before resuspension in RNase-free water. RNA concentration was determined using RNA ScreenTape Assay (Agilent Technologies Inc., Santa Clara, CA, USA) or by using Qubit RNA BR Assay Kit (ThermoFisher Scientific), both according to the manufacturer’s instructions.

Both RNA and protein amounts refer to batch culture measurements from a “standard culture volume”, e.g., 1 ml of culture normalized to an OD_600_ of 1.

### Protein extraction

For each condition, the volume equivalent to an OD_600_ of 3-5 per ml of a mid-exponentially growing culture was pelleted in a screw cap centrifuge tube, washed once in 1X PBS, flash frozen in liquid nitrogen and stored at -80 °C. Samples were thawed in 500 µl of 1X PBS containing protease inhibitor cocktail cOmplete^TM^ Ultra (Roche, Basel, Switzerland). Cells were disrupted as for the RNA extraction. Total protein content was determined using the BC Assay Protein Quantitation Kit (Interchim, Montluçon, France) according to the manufacturer’s instructions.

### Translation elongation rate assay

Translation elongation rates were measured via a fluorescent assay. *C. glutamicum* MB001(DE3) carrying the expression plasmid pMKEx2-*eyfp* [70] was cultivated in different media to yield a range of growth rates in a BioLector microcultivation system. One hour before induction of *eyfp* expression with isopropyl-β-D-thiogalactoside (IPTG), the BioLector 48-well flower plate was transferred to another BioLector instrument integrated into a robotics platform [78] and incubated under the same conditions. At mid-exponential phase *eyfp* expression was induced with 5.5 mM IPTG in triplicate. Immediately after induction, translation was stopped by adding 0.9 mg/ml chloramphenicol at 20-60 s intervals for a total of 15 time points. The robotics platform was used as it allows accurate time intervals for both the addition of IPTG and chloramphenicol. In control wells, IPTG, chloramphenicol or both compounds were substituted by the addition of an equivalent volume of cultivation medium. The BioLector flower plate was further incubated at 30 °C ON to allow full maturation of the EYFP fluorescent proteins. Cell growth (backscatter at 620 nm) and fluorescence (excitation 508 nm, emission 532 nm) were monitored every 5 minutes. After background correction, the specific fluorescence for each well [(fluorescence*ti* - fluorescence*t0*) / (backscatter*ti* - backscatter*t0*)] was calculated for each time point. The average specific fluorescence signal of a 3 h time interval after fluorescence reached a constant maximum was used to create the time-dependent translation plots. A linear regression was fit to the data and used to determine the time point at which the first fluorescent molecules appeared (y = 0). The translation elongation was calculated by dividing the number of amino acids in the EYFP molecule (238) by the determined time at which the first molecules appear.

### Self-replicator model and model calibration

The mechanistic coarse-grained modelling approach used in this study follows the work of Bosdriesz *et al*. [50]. The ODE-based self-replicator model sheds light on the connection of bacterial growth with intracellular nutritional states and associated governing rate parameters as consequence of protein synthesis and on growth-dependent ribosome synthesis regulation by means of a ppGpp-based mechanism. Herein, ribosomes forming a complex with unloaded tRNA, i.e. inactive non-translating ribosomes, provide a measure of ribosome inefficiency. The model considers amino acid biosynthesis generation, protein elongation, ppGpp-related metabolism and ribosome synthesis assuming a constant total proteome size and cell volume. The model was implemented in Matlab (Mathworks, Natick, MA, USA) and is provided in the Supplementary Data.

Organism-specific model parameters of key interest in this study are the maximum translation rate *k_R,max_* and the ribosome production rate *v_rrn_* (referred to as the *rrn* operon transcription initiation rate in [50]). After calibrating the qualitative nutrient quality (*k_n_*) for each of the studied steady-state growth conditions for *C. glutamicum*, rate parameters *k_R,max_* and *v_rrn_* were estimated from *µ* and *Φ_R_* (fraction of ribosomal proteins) measurements, while the remaining model parameters were set to literature values (see Supplementary Note 5, Table S6). For model calibration, 10 data points were available (Table S2). Each data point consists of the fraction of ribosomal protein (*Φ_R_*), derived from the fraction of total RNA per total protein, and the specific steady-state growth rate (*µ*), ranging from 0.07 h^-1^ to 0.94 h^-1^, at which *ΦR* was measured. Standard deviations for the measurements were determined for each data point by error propagation. Analogously, the model calibration procedure was performed for *E. coli* grown at 30 °C and for *E. coli* grown at 37 °C, based on publicly available data (see Supplementary Note 5C for details).

For parameter estimation, nonlinear weighted least-squares regression was applied. In short, the weighted residual sum-squared error between the model predicted and measured data were minimized based on a global gradient-free optimization heuristic. Differential equation systems were solved with Matlab’s ode15s solver. Standard deviations of the model parameters were calculated by parametric bootstrapping (500 runs with random initial parameter values). The *χ*^2^-test was applied to assess fit quality. All calculations were performed with custom Matlab scripts. Further details on the model, data processing, and calibration procedures are provided in the Supplementary Note 5. The resulting parameter settings of the three calibrated models used in this work are listed in Supplementary Tables S6 and S7.

To investigate the response in intracellular ribosome production to an upshift of the growth rate in *C. glutamicum* the medium was switched from comparably poor media (CGXII + glutamate, CGXII + ethanol) to a comparably rich medium (CGXII + glucose). To this end, steady-state growth rates *µ* and ribosomal protein fractions *Φ_R_* were simulated with the calibrated *C. glutamicum* model for each of the two poor media, mimicked via appropriate *k_n_* values (Supplementary Table S9). The nutrient upshift was simulated as follows: first, nutrient-poor conditions were simulated. The resulting steady-state values were then used as initial conditions for simulating the nutrient-rich condition. For maximal growth rate prediction (^μ̂^_max_), the upper limit of the maximum specific growth rates for *C. glutamicum* and *E. coli*, were extrapolated with the calibrated models by assuming a hypothetical nutritional quality value (*k_n_*) two orders of magnitude larger than the *k_n_* used for model calibration (Supplementary Table S8), while using the previously estimated *k_R,max_* and *v_rrn_* values.

### Western blotting

Samples from mid-exponential *C. glutamicum* wt or SM34 cultures grown in BHI + 2% (w/v) glucose were lysed by bead beating (Precellys24, Peqlab Biotechnologie, Erlangen, Germany) and cleared by centrifugation at 4 °C, 13000 rpm for 10 min. The supernantant was collected and protein concentration determined by BC Assay Protein Quantitation Kit (Interchim, Montluçon, France). Protein samples (20 µg) were separated by SDS-PAGE (Mini-protean TGX any-kD, Bio-Rad, Hercules, CA, USA) and transferred onto a PVDF membrane (0.2 µm, Novex, Thermo Fisher Scientific Inc., Waltham, MA, USA). Membranes were blocked with 5% (w/v) non-fat dried milk in PBS containing 0.1% (v/v) Tween20 (PBST) for 30 min at room temperature. The membranes were then incubated with either primary monoclonal antibodies anti-mCherry (Clontech #632543, Takara Bio USA, Inc., Mountain View, CA, USA) or anti-GFP (#CAU20008, Biomatik Corporation, Cambridge, Canada) diluted 1:2000 and 1:3000 in PBST, respectively. After washing with PBST, the membranes were incubated with the anti-mouse secondary antibody conjugated to alkaline phosphatase. Chemiluminescent signal detection was performed on a Fujifilm LAS-3000 imager (Fujifilm, Minato, Japan) after incubation with CDP-Star (Thermo Fisher Scientific Inc., Waltham, MA, USA).

## Data availability

The SMLM raw data used to generate Fig. 2b, 2c, 3 and Table S1 is provided as Source Data file. All other data can be obtained from the corresponding authors upon reasonable request.

## Author contributions

S.M. and M.B. designed the study. S.M., T.G., I.A. and J.H. performed the microscopy imaging acquisition. C.C.S. and K.N. developed the super-resolution emitter counter tool, SurEmCo. S.M., M.G. and R.T. contributed to the chemostat cultivations. S.M. and L.H. performed total RNA and protein extractions. S.M., N.T. and S.N. contributed to the translation elongation rate measurements. M.C. and K.N. adapted the self-replicator model and performed the simulations. S.M. performed the nutrient up-shit experiments. S.M., T.G., M.C., K.N. and M.B. analysed the data and wrote the manuscript as well as the Supplementary Information.

## Acknowledgements

The authors thank Professor Wolfgang Wiechert (Forschungszentrum Jülich) for fruitful discussions concerning the coarse-grained modeling effort as well as Professor Julia Frunzke (Forschungszentrum Jülich) for providing the pEKEx2-*eyfp* plasmid. This work was supported by the German Federal Ministry of Education and Research (BMBF; Grant 031A302C) and the CLIB-Competence Center Biotechnology (CKB) funded by the European Regional Development Fund ERDF (grant number 34.EFRE-0300097).

## Notes

### Competing Interest Statement

The authors have declared no competing interest.

