## Supplementrary notes, tables and figures for "Growth-rate dependency of ribosome abundance and translation elongation rate in *Corynebacterium glutamicum* differs from *Escherichia coli*"

### Supplementary files

#### Table of Contents

Supplementary Notes ... 2

Supplementary Note 1. SMLM ribosome counting with SurEmCo ... 2

Supplementary Note 2. Estimation of the number of ribosomes per cell from the R/P ratio values ... 5

Supplementary Note 3. Determination of the translation elongation rate via a fluorescent assay ... 6

Supplementary Note 4. Determination of the active ribosome fraction ... 7

Supplementary Note 5. Modelling the Rb/µ correlation for *C. glutamicum* ... 8

A. Preparing experimental data for modelling ... 8

B. Model overview and basic assumptions ... 9

C. Model calibration procedure ... 11

D. Discussion of the ribosome production rates in *C. glutamicum* ... 12

Supplementary Note 6. Comparison of modelling results for *C. glutamicum* and *E. coli* ... 14

Supplementary Tables ... 16

Table S1. SMLM data for ribosome quantification in *C. glutamicum* cells grown at different growth rates. ... 16

Table S2. RNA-protein (R/P) ratio of *C. glutamicum* at different growth rates. ... 17

Table S3. Translation elongation rate of *C. glutamicum* at different growth rates. ... 18

Table S4. Fraction of active ribosomes of *C. glutamicum* at different growth rates. ... 19

Table S5. Number of ribosomes per cell for *C. glutamicum* wt calculated from the R/P values. ... 20

Table S6. Model parameters and variables. ... 21

Table S7. Estimated model parameters. ... 23

Table S8. Prediction of maximum specific growth rate and ribosome protein fractions for high hypothetical nutrient qualities. ... 24

Table S9. Data for estimation of model parameters $k_{R,max}$ and $v_{rrn,max}$ used for nutrient upshift simulations. ... 25

Supplementary Figures ... 26

Figure S1. uS2-PAmCherry and bL19-EYFP protein fusions are produced in strain SM34. ... 26

Figure S2. Ribosome counting with the SurEmCo software. ... 28

Figure S3. Fluorescent assay for determination of the translation elongation rate. ... 29

Figure S4. Schematic of the coarse-grained self-replicator model and its most important functions. ... 30

Figure S5. Result of model calibration for *C. glutamicum* and *E. coli*. ... 31

Figure S6. Comparison of the Rb/µ correlation, inferred rates and active ribosome fraction observed in *C. glutamicum* and *E. coli* ... 32

Figure S7. Model-inferred tRNA loading status, ribosome activity and ppGpp concentrations across growth rates for *C. glutamicum* and *E. coli*. ... 34

Figure S8. Simulation of the upshift experiment – interrogation of the model. ... 36

Supplementary references ... 37

#### Supplementary Notes

##### Supplementary Note 1. SMLM ribosome counting with SurEmCo

The super-resolution emitter counter software SurEmCo (**https://github.com/modsim/suremco**) is developed in Python 3, using the numpy [1], scipy [2], numexpr, pandas [3] and OpenCV [4] libraries for data IO and processing. SurEmCo features an easy-to-use GUI (graphical user interface) provided by Qt (using PySide) and the VisPy library for 3D visualizations. Starting point for the analysis is a transmission image along with emitter positions previously reconstructed using SNSMIL. Cell regions are identified within the image (Fig. S2a): the image is background corrected (by division with a Gaussian blurred version of itself), inverted, rescaled, and subsequently binarized using a local thresholding algorithm [5]. Cell regions smaller than 0.42 $\mu m^2$ and larger than 42.25 $\mu m^2$ were discarded, yielding a separation of cells from background. Cell clusters and superposed cells were manually identified and excluded from the downstream analysis. Cell lengths and widths are approximated by fitting ellipses to the cell contour. For each detected cell, emitters are collected and per-cell emitter tracking is performed (Fig. S2b and S2c). Out of the different tracking algorithms available in SurEmCo, for performance reasons all images were analysed using a custom tracker written in C++, which implements an accelerated search by KD-trees [6] using the nanoflann library (**https://github.com/jlblancoc/nanoflann**).

In our study, emitter tracking is eased because the samples are fixed. Positional drifts in the obtained emitter traces may therefore only originate from movement of the sample stage, imperfect immobilization of the cells on the cover glass, residual mobility of the ribosomal proteins in the fixed cell, and/or imprecise position determination in the single molecule localization measurements.

In emitter tracking, two potential SMLM issues were tackled that typically arise in the determination of the number of single molecules per cell, namely (1) the presence of the same fluorophore in consecutive frames and (2) the blinking probability of the fluorophore used. During SMLM, the fluorescence of all single molecules was collected until they permanently photobleached. Under the applied illumination conditions (561 nm light power) the time until permanent photobleaching is ~50-1000 ms, i.e. a single photoactivated PAmCherry molecule is detectable in several consecutive images. The SurEmCo tracking algorithm that generates single-molecule time traces, checks for every detected emitter, whether there is an emitter in the previous image at that position, and counted every non-interrupted trace as one ribosomal protein (Fig. S2c and S2d emitters #1 and #2). The second important aspect considered is the blinking probability of the fluorophore. The fluorescence output of most fluorophores is not constant, but shows intensity fluctuations with prominent periods of emission (ON-periods) interrupted by periods of no emission (OFF-periods), although the excitation is kept unchanged. This behaviour cannot be observed in ensemble measurements since it is averaged out. However, in studies on single fluorophores, where the molecules were immobilized and/or observed over long periods of time it becomes apparent that it is a common behaviour (see for instance [7, 8]). When counting molecules, blinking becomes a problem since it introduces uncertainty on whether a new emerging photoactivated PAmCherry molecule is the result of a first-time photoactivation event or a molecule that returned from an interim OFF-state to an ON-state. To minimize the influence of the latter on the final emitter numbers by falsely counting returning events possibly multiple times, the spatio-temporal relationship of the emerging tracks is analysed. If an emitter is detected at the position of a terminated track within one image frame after the termination, the absence is regarded as a blinking event, meaning it is considered to be the same emitter and part of the former track (Fig. S2d emitter #3). This way the vast majority of the PAmCherry blinking events are accounted for. The detected emitters are displayed in a tabulated form (Fig. S2e) for further processing.

**Supplementary Note 2. Estimation of the number of ribosomes per cell from the R/P ratio values**

The RNA/protein ratio is a well described proxy of the ribosome abundance in the cell. In exponentially growing cells it is estimated that ~86% of total RNA is rRNA [9-11] and therefore the number of ribosomes in a sample can be calculated by $N_R = 0.86 \cdot R/m_{rRNA}$, where $N_R$ is the total ribosome number, $R$ is the total RNA content (here in 1 ml culture of $OD_{600} = 1$) and $m_{rRNA}$ is the mass of rRNA per ribosome. In *C. glutamicum* all 6 *rrn* operons are very similar in sequence [12, 13]; therefore, to calculate the molecular mass of a single ribosome the 5S, 16S and 23S rRNA sequences from the *rrnA* operon were used. They encompass 4,769 nucleotides with a total molecular mass of 1,522.98 kDa or $2.53 \times 10^{-15}$ mg. Results are shown in Table S5.

From experimentally derived ribosome numbers the *rrn* transcription initiation rate can be obtained by multiplication with the corresponding specific growth rate as $v_{rrn} = N_R \cdot \mu$ [10].

**Supplementary Note 3. Determination of the translation elongation rate via a fluorescent assay**

The *C. glutamicum* MB001(DE3) strain carrying the reporter plasmid pMKEx2-*eyfp* was used for all translation assays. The strain was pre-cultivated ON in the same medium as in the assay. A BioLector cultivation system was used for automated recording of backscatter as measure of growth and of fluorescence as measure of translation. At mid-exponential growth phase, expression of *eyfp* was induced with 5.5 mM IPTG and translation stopped by addition of 0.9 mg/ml chloramphenicol at 20-60 s intervals using a robotics platform to ensure accurate and reproducible intervals between experiments (Fig. S3a). The specific fluorescence was then calculated from the backscatter and fluorescence values for each time point measured (every 5 min). After normalization with the no-IPTG control, the mean specific fluorescence of a 3 h interval after the fluorescent signal reached a constant maximum was then plotted for each translation arrest time point. Finally, a linear regression was used to determine the intersection point on the *x*-axis (Fig. S3b). Calculation of the translation elongation rate took into account the 10 s determined by Dai *et al.* for the initial translation initiation steps in *E. coli* [14]. In our robotics setup there is a ~7 s delay between IPTG induction and chloramphenicol addition. Therefore the translation elongation rate was calculated as the ratio between the length of the EYFP protein (238 aa) and the time at which fluorescence first appears from which 3 s were deducted. Results for all tested conditions are shown in Fig. S3b and S3c.

**Supplementary Note 4. Determination of the active ribosome fraction**

The active ribosome fraction was calculated according to Dai *et al.* [14],

$$f_{active} = \frac{N_{activeRb}}{N_{Rb}} = \mu \frac{\sigma\prime}{k_R(\frac{R}{P})}$$

where $\sigma'$ is a dimensionless constant given by $m_{rRNA}/(0.86 \cdot m_{aa})$. For *C. glutamicum* the $m_{rRNA}$ is 1522.98 kDa (see Supplementary Note 2). The average molecular weight of an amino acid ($m_{aa}$) is 110 Da, thus for *C. glutamicum* $\sigma' \approx 1.61 \times 10^4$. The active ribosome fraction was then calculated from the values for growth rate ($\mu$) and R/P ratio listed in Table S4 and the translation elongation rate ($k_R$) values from Table S3. The difference in $\mu$ values between Tables S3 and S4 comes from the fact that due to the experimental setup the same cultures could not be used for both the R/P ratio measurements and the translation elongation assays. Therefore, the mean $\mu$ between the biological replicates for the two assays was determined and used in the calculation of the active ribosome fraction.

**Supplementary Note 5. Modelling the Rb/μ correlation for *C. glutamicum***

Here we describe preparation of experimental data prior to modelling and the essential assumptions of the self-replicator model. In addition, further details about model calibration and results from simulations are provided.

**A. Preparing experimental data for modelling**

*Calculation of ribosomal protein fractions $\Phi_R$:* Fractions of total ribosomal protein per total protein (w/w) ($\Phi_R$) were derived from measurements of total RNA per total protein (w/w) determined under steady-state conditions according to the procedure described in Scott *et al.* [15]. In short, $\Phi_R$ for a specific growth rate was obtained by multiplying RNA/protein ratios by a factor $\rho$ [extended ribosome (μg)/RNA (μg)] [15]. As described in Scott *et al.*, the extended ribosome encompasses all ribosomal proteins plus their affiliates which include all initiation and elongation factors as well as tRNA synthases, etc. $\rho$ is calculated as the product of the three ratios: a) rRNA per total RNA (~86% w $w^{-1}$), b) ribosomal protein per rRNA (~53% w $w^{-1}$), and c) affiliated ribosomal proteins per ribosomal protein (~167% w $w^{-1}$). For *E. coli*, the value of $\rho$ was estimated to be ~0.76 [15]. Due to the lack of organism-specific data for *C. glutamicum*, the same value was also used in this study.

*Determination of specific growth rates $\mu$:* Specific growth rates $\mu$ ($h^{-1}$) were calculated from doubling times (min) by means of $\mu = \ln(2)/(\textit{doubling time}/60)$ or from doubling rates (doublings $h^{-1}$) by means of $\mu = \textit{doubling rate} \cdot \ln(2)$.

#### B. Model overview and basic assumptions

A schematic of the most important model components and variables is shown in Fig. S4. For the full set of model equations, the reader is referred to the Supplementary model files and the original study [16]. A listing of model parameters is found in Table S6. Here, only most relevant modelling assumptions are given:

- The specific growth rate $\mu$ ($h^{-1}$) is defined as the instantaneous volumetric translation elongation rate $v_R$ ($\mu mol_{aa}\ L_{cell}^{-1}\ s^{-1}$) per total amino acid concentration (total proteome) $p$ ($\mu M_{aa,tot}$) of the cell (Eq. 1). $v_R$ is determined by the maximum specific translation rate $k_{R,max}$, which is an independent parameter, and instantaneous concentrations of ribosomal proteins $r$ (µM), amino-acylated tRNA $t_{aa}$ (µM) and free tRNA $t_f$ (µM). The instantaneous specific translation elongation rate $k_R$ (aa ribosome$^{-1}$ s$^{-1}$) was calculated from dividing $v_R$ by $r$ at a given nutrient condition. While $v_R$, $k_R$, $t_{aa}$, $t_f$ and $r$ are model variables, the total proteome $p$ is assumed to be constant and $k_R$ a free parameter.
- The availability of the amino acids required for translation is reflected by the maximal specific rate of amino acid synthesis, referred to as nutrient quality parameter $k_n$ ($s^{-1}$), which is (due to coarse-graining) an aggregated parameter that simultaneously captures the catalytic efficiency, extracellular nutrient quality, and nutrient concentrations.
- Ribosomal proteins are produced at an instantaneous rate $v_{rrn}$ (ribosomes $s^{-1}$), which is equivalent to (*rrn* initiations $s^{-1}$ cell$^{-1}$), that is derived from the maximum ribosome production rate $v_{rrn,max}$ and the instantaneous ppGpp concentration ppGpp (µM) (Eq. 2). Herein, $v_{rrn,max}$ is an independent parameter.

- The fraction of ribosomal proteins $\Phi_R$ is determined by the concentration of all amino acids within the ribosomal proteins, calculated as $r{\cdot}N_{a,r}$ (µM) per total proteome $p$ (Eq. 3), where $N_{a,r}$ is the number of amino acids per extended ribosome (12307).
- Three different ribosome species are considered (Eq. 4):

  (1) Ribosomes that form complexes with amino acid-loaded tRNA ($r_{taa}$).

  (2) Ribosomes that form complexes with unloaded amino acid tRNA ($r_{tf}$).

  (3) Free ribosomes that are not associated with any tRNA molecule ($r_f$).

  While the first ribosome species is considered to actively translating, the latter two species do not contribute to the instantaneous translation elongation rate. Consequently, it is straightforward to calculate the fraction of actively translating ribosomes ($f_{active}$) (Eq. 5).

$$\mu = \frac{v_R}{p} = \frac{k_{R,max} \cdot r \cdot f(t_{aa}, t_f, \dots)}{p} \quad \text{(Eq. 1)}$$

$$v_{rrn} = v_{rrn,max} \cdot \frac{RNAP_F}{K_{M,rrn} + RNAP_F} \cdot \frac{1}{1 + \frac{ppGpp}{k_{i,ppGpp}}} \quad \text{(Eq. 2)}$$

$$\Phi_R = (r/p) \cdot N_{a,r} \quad \text{(Eq. 3)}$$

$$r = r_{taa} + r_{tf} + r_f \quad \text{(Eq. 4)}$$

$$f_{active} = \frac{r_{taa}}{r_{taa} + r_{tf} + r_f} \quad \text{(Eq. 5)}$$

#### C. Model calibration procedure

The model calibration procedure was performed for *C. glutamicum*, for *E. coli* grown at 30 °C, and for *E. coli* grown at 37 °C, independently, using appropriate data sets. Data for *C. glutamicum* were produced in this work (Tables S2 and S9). Data for *E. coli* were taken from the literature, specifically from [17] and from [10, 18] for 30 °C and 37 °C, respectively. This amounted to a total of 19 (30 °C) and 33 (37 °C) data points. In both cases, measurement errors of 10-15% were assumed according to values reported in [15]. The three data sets can be found in Supplementary Table S2 and in the Supplementary Model File.

Appropriate nutrient quality values $k_n$ for each of the available growth conditions were found by means of the self-replicator model (Supplementary Note 5B). Here, $k_n$ is the effective maximal rate of amino acid synthesis per unit of metabolic enzyme that summarizes the catalytic efficiency of the overall cellular metabolism, the extracellular nutrient quality, nutrient concentration and related factors. Measurements for $\mu$ and $\Phi_R$ and the corresponding $k_n$ values are listed in Table S9. Upper and lower values for $k_n$ were taken from [16]. Then, the key parameters, namely the maximum translation rate $k_{R,max}$ and the maximum ribosome production rate $v_{rrn,max}$, were estimated with the coarse-grained model from these sets of ($\mu$, $\Phi_R$, $k_n$)-triplets as described in the Materials and Methods. Measured and simulated data agreed very well for all strains and conditions (Fig. 4a). The sum of squared residuals (SSR) were ~22, 29, and 33 for *C. glutamicum*, *E. coli* 30 °C and *E. coli* 37 °C, respectively. These values were below the acceptable chi-squared test statistic of ~29, 51 and 84 at 95% significance level, respectively. Best parameter estimates for $k_{R,max}$ and $v_{rrn,max}$ including their associated standard deviations are found in Table S7 for *C. glutamicum* and *E. coli*. The resulting model fits are shown in Fig. S5.

#### D. Discussion of the ribosome production rates in *C. glutamicum*

In the model $v_{rrn}$ is determined from $v_{rrn,max}$ (Table S7) and other parameters such as the parameter for free RNA polymerase concentration ($RNAP_f$) (Table S6) by means of a pseudo kinetic (see (Eq. 2) in Supplementary Note 5B). Therefore, we only discuss how $v_{rrn,max}$ is qualitatively interrelated with the instantaneous ribosome production rate $v_{rrn}$. A good match was found between the model-inferred instantaneous ribosome production rate $v_{rrn}$ and the rates independently approximated from the SMLM counted ribosome numbers (Fig. S6c, Supplementary Note 2). Model-based inference shows that the ribosome production rate is ~3 newly extended ribosomes (*rrn* operon initiations) per second and cell at $\mu$ = 0.4 $h^{-1}$ (Fig. S6c). At the predicted maximal growth rate of $\widehat{\mu}_{max}$ = 0.94 $h^{-1}$, enabled by using a hypothetical "super-rich" medium, an instantaneous ribosome production rate of ~11 *rrn* initiations $s^{-1}$ $cell^{-1}$ was estimated. To summarize, values of $v_{rrn,max}$ (Table S7) exceed those of the corresponding instantaneous rates $v_{rrn}$ (Fig. S6c, dashed lines) by 1-2 orders of magnitude, which we attribute to the pseudo kinetics and their parameters, except $v_{rrn}$ as explained before.

To unravel what ultimately limits ribosome production to achieve even higher growth rates > $\mu_{max}$, it should be noted, that the concentration of unloaded tRNA (Fig. S7b) and, therewith, the concentration of non-translating ribosome complexes drops towards zero (Fig. S7d). Modelling results reveal that, at higher growth rates, the insignificant number of the remaining non-translating ribosome complexes is connected to a very low ppGpp concentration (Fig. S7e) limiting the further increase in ribosomes and therewith settle the maximum growth rate.

**Supplementary Note 6. Comparison of modelling results for *C. glutamicum* and *E. coli***

Comparison between modelling results for *C. glutamicum* grown at 30 °C and *E. coli* grown at 30 °C and 37 °C revealed (after appropriate model calibration with literature data, see Materials and Methods, Supplementary Note 5C, and Supplementary Table S6) that *C. glutamicum* and *E. coli* at 30 °C require similar ribosome abundance to support comparable growth rates. These values are considerably higher compared to *E. coli* at 37 °C for growth rates $\mu$ = 0.4-1.2 $h^{-1}$ (Fig. S6a). For *E. coli*, the inferred maximum translation elongation rates were $k_{R,max}$ = 8.3 ± 0.6 aa $ribosome^{-1}$ $s^{-1}$ and 17.7 ± 0.6 aa $ribosome^{-1}$ $s^{-1}$ grown at 30 °C and 37 °C, respectively (Table S7, Fig. S6b), which is in line with previous experimental results [14, 19]. *rrn* initiation rates were inferred in the range $v_{rrn}$ = 1.1 ± 0.4 – 18.4 ± 6.5 *rrn* initiations $s^{-1}$ $cell^{-1}$ and 2.4 ± 0.3 – 20.0 ± 2.5 *rrn* initiations $s^{-1}$ $cell^{-1}$ for *E. coli* at 30 °C and 37 °C, respectively. While values of $k_{R,max}$ were comparable for *C. glutamicum* and *E. coli* (30 °C), they only exhibited ~50% of the value obtained for *E. coli* grown at 37 °C. Similarly, at $\mu$ = 0.9 $h^{-1}$ *C. glutamicum* and *E. coli* at 30 °C exhibit ~2-fold increased ribosome production rates compared to *E. coli* at 37 °C, to meet the increased demand for ribosomes and to compensate for the lower translation rates (Fig. S6c and Supplementary Note 5D).

The order of magnitude of model-derived ribosome production rates could be experimentally verified for *E. coli* if we assume that 55,000 ribosomes at $\mu$ = 0.77 $h^{-1}$ (30 °C) [20] and 72,000 ribosomes at $\mu$ = 1.72 $h^{-1}$ (37 °C)[10] yield $v_{rrn}$ = 9.5 and 34.4 *rrn* initiations $s^{-1}$ $cell^{-1}$, respectively (see Supplementary Note 2 and Fig. S6c). In line with the findings for *C. glutamicum* all values of $v_{rrn,max}$ (Table S7) exceed those of the corresponding instantaneous rates $v_{rrn}$ (Fig. S6c, dashed lines) by 1-2 orders of magnitude. Compared to *C. glutamicum*, maximum ribosome production rates $v_{rrn,max}$ were found to be 2.5- and 3.7-fold increased for *E. coli* grown at 30 °C and 37 °C, respectively. Consistent with this, also the instantaneous ribosome production rates $v_{rrn}$ obtained at large $k_n$ values (super-rich medium, Table S8) were found to be 2.1 and 2.6 fold increased for *E. coli* grown at 30 °C and 37 °C, respectively, when compared to *C. glutamicum*.

By increasing the specific rate of amino acid production ($k_n$, Table S6), maximum specific growth rates $\hat{\mu}_{max}$ = 0.94, 1.31 and 2.03 $h^{-1}$ were predicted for *C. glutamicum*, *E. coli* at 30 °C and *E. coli* at 37 °C, respectively (see Materials and Methods, Table S8, Fig. S6a-d). Notably, the predicted maximal growth rates for *E. coli* are close to the maximum growth rates $\mu$ = 1.26 $h^{-1}$ (30 °C) and 2.3 $h^{-1}$ (37 °C) that were previously experimentally observed [17, 21]. For *C. glutamicum*, as limiting factor for ribosome production (Supplementary Note 5D) and therefore higher growth rates, the model points to the limited availability of spare tRNAs (Fig. S7b). As growth rate increases, in *C. glutamicum* the concentration of unloaded tRNA (Fig. S7b) and, therewith, the concentration of non-translating ribosome complexes drops towards zero more rapidly than in *E. coli* (Fig. S7d) and consequently settles a lower maximum growth rate. Because *E. coli* is known to employ active mechanisms of ribosome deactivation at growth rates $\mu < 0.5$ $h^{-1}$ to maintain a relatively high translation rate at 37 °C (Fig. S6b, [14]), a direct comparison of how *E. coli* and *C. glutamicum* utilize ribosomes at low growth rates is not possible with our model. Therefore, regarding *E. coli* grown at 37 °C, the model was only calibrated with experimental data in the range $\mu \geq 0.5$ $h^{-1}$.

#### Supplementary Tables

**Table S1. SMLM data for ribosome quantification in *C. glutamicum* cells grown at different growth rates.** *C. glutamicum* dual labelled ribosome strains (SM34 and SM55) were grown under different nutrient/cultivation conditions to obtain a range of growth rates. Cell volume was calculated for each individual cell following the formula: $V = \pi W^2(L - W/3)/4$ [22]. For each condition, at least 3 independent biological replicates were analysed, except for the chemostat and SM55 experiments where a minimum of 2 technical replicates were analysed. Shown are the mean and standard deviation for each of the parameters measured.

| Strain/Cultivation | Growth rate ($\mu$) ($h^{-1}$) | n | Ribosomes/cell | Length ($L$) (µm) | Width ($W$) (µm) | Volume ($V$) ($\mu m^3$) | Ribosomes/$\mu m^3$ |
|---|---|---|---|---|---|---|---|
| SM34_CGXII+PYR (EXP) | 0.31±0.02 | 179 | 14,474±5,097 | 1.73±0.35 | 0.89±0.10 | 0.92±0.36 | 16,431±4386 |
| SM34_CGXII+PYR+CAA (EXP) | 0.40±0.02 | 110 | 22,020±8,221 | 2.09±0.38 | 0.92±0.09 | 1.21±0.41 | 18,695±4,953 |
| SM34_CGXII+GLU (EXP) | 0.43±0.01 | 281 | 25,770±9,508 | 1.98±0.40 | 0.92±0.11 | 1.17±0.50 | 24,046±8,326 |
| SM34_CGXII+GLU+CAA (EXP) | 0.46±0.03 | 239 | 34514±11,111 | 2.29±0.48 | 0.96±0.13 | 1.48±0.64 | 26,059±10,712 |
| SM34_BHI+GLU (EXP) | 0.47±0.03 | 123 | 45,207±25,667 | 2.26±0.73 | 1.02±0.15 | 1.67±1.14 | 29,241±9,630 |
| SM34_BHI+GLU+CAA (EXP) | 0.48±0.03 | 98 | 60,680±19,698 | 2.71±0.67 | 1.07±0.12 | 2.19±0.92 | 29,751±7,961 |
| SM34_CGXII+PYR (STA) | | 113 | 13,351±4,750 | 1.78±0.35 | 0.90±0.09 | 0.95±0.33 | 14,725±4,534 |
| SM34_CGXII+PYR+CAA (STA) | | 158 | 15,361±5,977 | 1.84±0.37 | 0.89±0.10 | 0.99±0.41 | 16,074±4,559 |
| SM34_CGXII+GLU (STA) | | 264 | 19,629±11,376 | 2.04±0.50 | 1.01±0.16 | 1.44±0.72 | 14,312±5,615 |
| SM34_CGXII+GLU+CAA (STA) | | 266 | 30,690±10,086 | 2.44±0.56 | 1.08±0.12 | 1.98±0.86 | 16,415±3,783 |
| SM34_BHI+GLU (STA) | | 239 | 16,762±5,212 | 1.85±0.34 | 1.04±0.10 | 1.30±0.48 | 13,320±2817 |
| SM34_BHI+GLU+CAA (STA) | | 137 | 20,985±8,732 | 1.93±0.38 | 1.12±0.13 | 1.59±0.67 | 13,497±3,497 |
| SM34_Chemostat (0.20) | 0.2 | 30 | 11,942±3,084 | 1.74±0.27 | 0.88±0.06 | 0.89±0.21 | 13,888±3,945 |
| SM34_Chemostat (0.30) | 0.3 | 31 | 17,293±7,545 | 1.94±0.34 | 0.88±0.14 | 1.05±0.46 | 16,778±3,321 |
| SM34_Chemostat (0.43) | 0.43 | 47 | 33,859±15,211 | 2.39±0.47 | 0.98±0.07 | 1.59±0.48 | 21,266±6,568 |
| SM55_CGXII+PYR (EXP) | 0.24±0.00 | 34 | 13,790±7,171 | 1.98±0.34 | 0.96±0.12 | 1.24±0.49 | 11,082±3,477 |
| SM55_CGXII+GLU (EXP) | 0.41±0.02 | 86 | 27,723±12,820 | 1.98±0.47 | 1.01±0.11 | 1.37±0.59 | 21,165±7,440 |
| SM55_BHI+GLU (EXP) | 0.49±0.00 | 80 | 84,020±33,842 | 3.16±0.74 | 1.17±0.17 | 3.18±1.58 | 28,193±7,363 |

**Table S2. RNA-protein (R/P) ratio of *C. glutamicum* at different growth rates.** *C. glutamicum* ATCC 13032 strain (wt) or an evolved variant of this strain (EVO5) [23] were grown under different nutrient/cultivation conditions to obtain a range of growth rates. The R/P ratio data are shown in Fig. 3c and 4a. For each condition total RNA contents and total protein contents represent the results of at least two biological and two technical replicates. Shown are the mean and standard deviation for each of the parameters measured. Standard deviation for R/P ratio was calculated assuming error propagation.

| **Strain** | **Medium** | **Growth rate (*μ*) ($h^{-1}$)** | **Total RNA (μg $OD_{600}^{-1}$ $ml^{-1}$)** | **Total protein (μg $OD_{600}^{-1}$ $ml^{-1}$)** | **R/P ratio** |
|---|---|---|---|---|---|
| EVO5 | BHI+CGXII+GLU | 0.94±0.08 | 66.58±7.02 | 94.96±5.6 | 0.70±0.08 |
| EVO5 | BHI+GLU | 0.77±0.07 | 38.83±3.47 | 68.30±3.91 | 0.57±0.06 |
| wt | BHI+GLU | 0.61±0.02 | 30.81±2.24 | 95.10±10.20 | 0.32±0.04 |
| wt | CGXII+GLU | 0.47±0.03 | 23.71±6.33 | 97.58±11.83 | 0.24±0.07 |
| wt | CGXII+ACE | 0.40±0.01 | 14.43±1.51 | 71.01±2.89 | 0.20±0.02 |
| wt | CGXII+PYR | 0.29±0.01 | 14.76±1.40 | 78.06±0.83 | 0.19±0.02 |
| wt | CGXII+LAC | 0.27±0.07 | 13.50±1.11 | 86.54±4.47 | 0.16±0.02 |
| wt | CGXII+EtOH+CAA | 0.24±0.05 | 13.48±3.37 | 74.67±8.70 | 0.18±0.05 |
| wt | CGXII+EtOH | 0.12±0.01 | 11.28±0.88 | 79.51±14.08 | 0.14±0.03 |
| wt | CGXII+Glutamate | 0.07±0.02 | 11.81±2.13 | 82.03±8.55 | 0.14±0.03 |
| wt | BHI+GLU (STA) | 0 | 12.55±2.83 | 77.02±10.49 | 0.16±0.04 |

**Table S3. Translation elongation rate of *C. glutamicum* at different growth rates.** *C. glutamicum* MB001(DE3) carrying the reporter plasmid pMKEx2-*eyfp* was grown under different nutrient conditions to obtain a range of growth rates. The translation elongation results are shown in Fig. 4b. For each condition at least 2 biological and 3 technical replicates were assayed. Shown are the mean and standard deviation for each of the parameters measured.

| Strain | Medium | Growth rate ($\mu$) ($h^{-1}$) | Translation elongation rate ($k_R$) (aa $s^{-1}$) |
|---|---|---|---|
| MB001(DE3) pMKEx2-*eyfp* | BHI+CGXII+GLU | 0.79±0.14 | 9.00±0.13 |
| MB001(DE3) pMKEx2-*eyfp* | CGXII+GLU | 0.49±0.02 | 7.93±0.19 |
| MB001(DE3) pMKEx2-*eyfp* | CGXII+EtOH+CAA | 0.29±0.02 | 6.76±0.55 |
| MB001(DE3) pMKEx2-*eyfp* | CGXII+PYR | 0.20±0.02 | 5.29±0.16 |
| MB001(DE3) pMKEx2-*eyfp* | CGXII+EtOH | 0.18±0.03 | 4.91±0.17 |
| MB001(DE3) pMKEx2-*eyfp* | CGXII+LAC | 0.18±0.02 | 4.75±0.29 |
| MB001(DE3) pMKEx2-*eyfp* | CGXII+Glutamate | 0.07±0.00 | 1.89±0.06 |
| MB001(DE3) pMKEx2-*eyfp* | BHI+CGXII+GLU (STA) | 0 | 0.80±0.11 |

**Table S4. Fraction of active ribosomes of *C. glutamicum* at different growth rates.** *C. glutamicum* MB001(DE3) carrying the reporter plasmid pMKEx2-*eyfp* was grown under different nutrient conditions to obtain a range of growth rates. The fraction of active ribosomes was calculated according to Dai *et al.* [14] [$f_{activeRb}=\mu \cdot \sigma' / k \cdot (R/P)$]. The R/P values were determined for MB001(DE3) pMKEx2-*eyfp* grown in the same conditions as for the translation elongation assay. Shown is the mean between the growth rate from the translation elongation assays and those for R/P determination. The active ribosome fraction results are shown in Fig. 4c. For each condition, at least 2 biological and 3 technical replicates were assayed. Shown are the mean and standard deviation for each of the parameters measured.

| Strain | Medium | R/P | Growth rate ($\mu$) ($h^{-1}$) | Active ribosome fraction |
|---|---|---|---|---|
| MB001(DE3) pMKEx2-*eyfp* | BHI+CGXII+GLU | 0.40±0.04 | 0.73±0.09 | 0.91±0.08 |
| MB001(DE3) pMKEx2-*eyfp* | CGXII+GLU | 0.30±0.06 | 0.48±0.01 | 0.90±0.17 |
| MB001(DE3) pMKEx2-*eyfp* | CGXII+EtOH | 0.19±0.02 | 0.18±0.01 | 0.85±0.09 |
| MB001(DE3) pMKEx2-*eyfp* | CGXII+Glutamate | 0.19±0.03 | 0.06±0.01 | 0.73±0.12 |

**Table S5. Number of ribosomes per cell for *C. glutamicum* wt calculated from the R/P values.** *C. glutamicum* wt was grown in either BHI+GLU or CGXII+GLU until mid-exponential phase. Mean colony forming units (cfu) per OD were determined by plate enumeration of serial dilutions for two biological replicates. Total number of ribosomes was calculated according to $N_{Rb}=0.86 \cdot R/m_{rRNA}$ using the total RNA (µg $OD_{600}^{-1}$ $ml^{-1}$) values shown in Table S2.

| Strain | Medium | cfu/$OD_{600}$ | Ribosome number/$OD_{600}$/ml | $N_{Rb}$/cell |
|---|---|---|---|---|
| wt | BHI+GLU | $2.39 \times 10^{8}$ | $1.05 \times 10^{13}$ | 43,821 |
| wt | CGXII+GLU | $3.56 \times 10^{8}$ | $8.06 \times 10^{12}$ | 22,646 |

**Table S6. Model parameters and variables.** If not stated differently, values were taken from Bosdriesz *et al.* [16] and assumed to be similar for *C. glutamicum*.

| Parameter | Value | Unit | Description |
|---|---|---|---|
| *Amino acid metabolism* | | | |
| $k_i$ | 100 | µM | Feedback inhibition constant for amino acid synthesis |
| $k_n$ | Tables S8-S9 | $s^{-1}$ | Maximum rate of amino acid synthesis, reflects "nutrient quality" |
| $S_{tot,i}$ | 1 | µM | Total concentration of aminoacyl-tRNA synthetase for $a_i$ |
| $k_{S,i}$ | 100 | $s^{-1}$ | $k_{cat}$ *value* of aminoacyl-tRNA synthetase |
| $K_{M,a,i}$ | 100 | µM | Michaelis constant of aminoacyl-tRNA synthetase for amino-acids |
| $K_{M,t,i}$ | 1 | µM | Michaelis constant of aminoacyl-tRNA synthetase for uncharged tRNA |
| *Protein elongation* | | | |
| $k_{ta}$ | 1 | µM | Dissociation-constant of charged tRNA-ribosome complex formation |
| $k_t$ | 500 | µM | Dissociation-constant of uncharged tRNA-ribosome complex |
| $k_{R,max}$ | Table S7 | aa $rib^{-1}$ $s^{-1}$ | Maximum specific translation rate |
| $t_{tot,i}$ | 0.5 r | µM | Total tRNA conjugate to $a_i$. tRNA is proportional to ribosomes, with $\sum t_{tot,i}/r$ = 10 |
| *ppGpp metabolism* | | | |
| $RelA_{tot}$ | 1/15 | µM | RelA concentration (100 molecules per cell) |
| $k_{RelA}$ | 75 | $s^{-1}$ | $k_{cat}$ value of *ppGpp* synthesis by RelA |
| $k_{DrelA}$ | 0.26 | µM | Concentration of uncharged tRNA that leads to half maximal ppGpp production (interpretation depends on mechanism of ppGpp synthesis assumed) |
| $V_{ppGpp,syn}$ | $10^{-3}$ | µM $s^{-1}$ | Rate of *ppGpp* synthesis |
| $K_{ppGpp,deg}$ | ln(2)/30 | $s^{-1}$ | Half-life time of *ppGpp* after upshift is approximately 30 s |
| *Ribosome synthesis* | | | |
| $k_{i,ppGpp}$ | 1 | µM | Inhibition constant of *rrn* synthesis for *ppGpp* |
| $K_{M,rrn}$ | 20 | µM | Michaelis constant of *rrn*-promotors for free RNA polymerase ($RNAP_F$) |
| $V_{rrn,max}$ | Table S7 | $s^{-1}$ | Maximum ribosome production rate (referred to as maximum *rrn* operon transcription initiation rate in [16]) |
| $RNAP_F$ | 1 | µM | $RNAP_F$ Concentration of free RNA polymerase |
| *Related cell parameters* | | | |
| $V_{cell}$ | $2.5\cdot10^{-9}$ | µl | Cell volume |
| $N_{a,cell}$ | $15\cdot10^{8}$ | - | Total number of amino acids per cell |
| $N_{a,r}$ | 12307 | - | Number of amino acids per extended ribosome, including affiliates such as elongation factors |
| $N_{a,m}$ | 300 | - | Average number of amino acids in a metabolic protein (BNID:100017) |
| $N_{AV}$ | $6.022\cdot10^{23}$ | $mol^{-1}$ | Avogadro number |
| $p$ | $10^{6}$ | µM | Calculated total amino acid concentration in the cell; referred to as total proteome: $N_{a,cell}/(V_{cell}\ N_{Av})$ |
| **Variable** | | **Unit** | **Description** |
| $\mu$ | | $s^{-1}$, $h^{-1}$ | Specific growth rate |

| | | |
|---|---|---|
| $v_R$ | $\mu M_{aa}\ s^{-1}$ | Volumetric translation rate |
| $k_R$ | aa $rib^{-1}\ s^{-1}$ | Instantaneous specific translation rate |
| $v_{rrn}$ | $s^{-1}$ | Instantaneous ribosome production rate, *rrn* operon transcription initiation rate |
| $r$ | µM | Total ribosome concentration |
| $r_{taa}$ | µM | Concentration of tRNA-(loaded with amino acids) ribosome complex |
| $r_{tf}$ | µM | Concentration of tRNA-(unloaded with amino acids) ribosome complex |
| $r_f$ | µM | Concentration of free ribosomes |
| $t_{aa}$ | µM | Concentration of tRNA, loaded with amino acids |
| $t_f$ | µM | Concentration of free tRNA |
| *ppGpp* | µM | Concentration of *ppGpp* |
| $\Phi_R$ | - | Fraction of ribosomal proteins |
| $f_{active}$ | - | Fraction of active ribosomes |

**Table S7. Estimated model parameters.** Both the maximum translation $k_{R,max}$ (aa ribosome$^{-1}$ s$^{-1}$) and maximum ribosome production rate $v_{rrn,max}$ (*rrn* operon transcription initiations s$^{-1}$ cell$^{-1}$) were jointly estimated from experimental data ($\mu$, $\Phi_R$) (Table S9). The estimation procedure is described in the Materials and Methods.

| Organism | $k_{R,max}$ | Range | $v_{rrn,max}$ | Range |
|---|---|---|---|---|
| *C. glutamicum* | 9.3 ± 0.8 | [5-15] | 369.9 ± 127.8 | [200-2000] |
| *E. coli*, 30 °C | 8.3 ± 0.6 | [8-15] | 928.7 ± 326.1 | [400-2000] |
| *E. coli*, 37 °C | 17.7 ± 0.6 | [4-20] | 1384.2 ± 171.1 | [200-2000] |

**Table S8. Prediction of maximum specific growth rate and ribosome protein fractions for high hypothetical nutrient qualities.** For predictions, the calibrated model with parameter values given in Table S6 was used and $k_n$ was increased by two orders of magnitude compared to values found for “normal” conditions (Table S9). Sensitivity analyses of $k_n$ revealed that the prediction of $\hat{\mu}_{max}$ is relatively insensitive to changes in values of $k_n$, in particular for $k_n > 30$ $s^{-1}$ (data not shown).

| **Organism** | **Predicted $\hat{\mu}_{max}$ ($h^{-1}$)** | **Predicted $\hat{\Phi}_{R,max}$** | **Hypothetical $k_n$ ($s^{-1}$)** |
|---|---|---|---|
| *C. glutamicum* | 0.94 | 0.37 | 30.0 |
| *E. coli*, 30 °C | 1.31 | 0.56 | 33.5 |
| *E. coli*, 37 °C | 2.03 | 0.42 | 33.5 |

**Table S9. Data for estimation of model parameters $k_{R,max}$ and $v_{rrn,max}$ used for nutrient upshift simulations.** Experimentally determined steady-state growth rates $\mu$ and ribosomal protein fractions $\Phi_R$ at different nutrient conditions $k_n$. Specific growth rates are the same as in Table S2. Values $\Phi_R$ were calculated from total RNA and total protein content according to Supplementary Note 5A. The calibrated qualitative nutrient quality values $k_n$ are listed for each growth condition. Values of $k_n$ were set within the range reported by Bosdriesz *et al.* [16]. For simulation of the upshift experiments, values of $k_n$ (right column) were chosen in such a way that the obtained simulated steady state growth rates closely matched the experimentally observed growth rates ($\mu$=0.06, 0.17 and 0.46 $h^{-1}$ for CGXII+glutamate-, EtOH- and GLU-medium, respectively; see also Fig. 5).

| **Strain** | **Medium** | **Growth rate ($\mu$) ($h^{-1}$)** | **Ribosomal protein fraction ($\Phi_R$)** | **Nutrient quality ($k_n$) ($s^{-1}$)** | **Upshift ($k_n$) ($s^{-1}$)** |
|---|---|---|---|---|---|
| EVO5 | BHI+CGXII+GLU | 0.94±0.08 | 0.53±0.06 | 0.240 | |
| EVO5 | BHI+GLU | 0.77±0.07 | 0.43±0.05 | 0.160 | |
| wt | BHI+GLU | 0.61±0.02 | 0.23±0.03 | 0.075 | |
| wt | CGXII+GLU | 0.47±0.03 | 0.18±0.05 | 0.049 | 0.051 |
| wt | CGXII+ACE | 0.40±0.01 | 0.15±0.02 | 0.041 | |
| wt | CGXII+PYR | 0.29±0.01 | 0.14±0.02 | 0.029 | |
| wt | CGXII+LAC | 0.27±0.07 | 0.12±0.02 | 0.027 | |
| wt | CGXII+EtOH+CAA | 0.24±0.05 | 0.14±0.04 | 0.023 | |
| wt | CGXII+EtOH | 0.12±0.01 | 0.11±0.02 | 0.012 | 0.016 |
| wt | CGXII+Glutamate | 0.06±0.02 | 0.11±0.02 | 0.006 | 0.006 |

#### Supplementary Figures

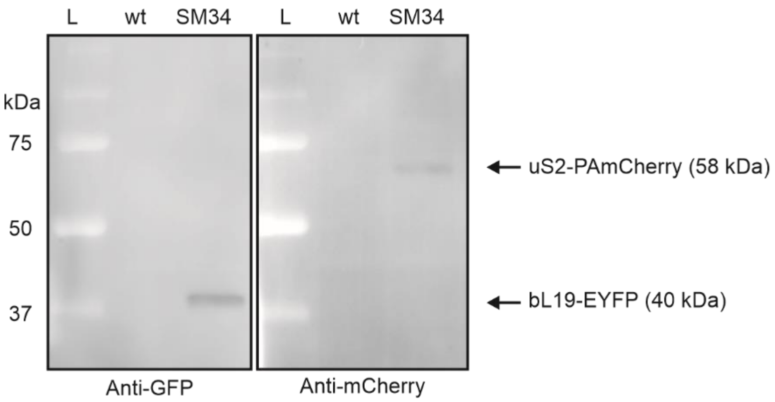


**Figure S1. uS2-PAmCherry and bL19-EYFP protein fusions are produced in strain SM34.** Cell lysates of mid-exponential *C. glutamicum* wt and SM34 cultivated in BHI + glucose were separated by SDS-PAGE and analysed by western blot using the indicated antibodies. On the left, the molecular mass (kDa) of relevant proteins from the Dual Color Precision Plus Protein Prestained Standards (Bio-Rad) on lanes L is shown. Arrows show the detection position of each fusion protein in the SM34 extracts along with their predicted molecular mass.

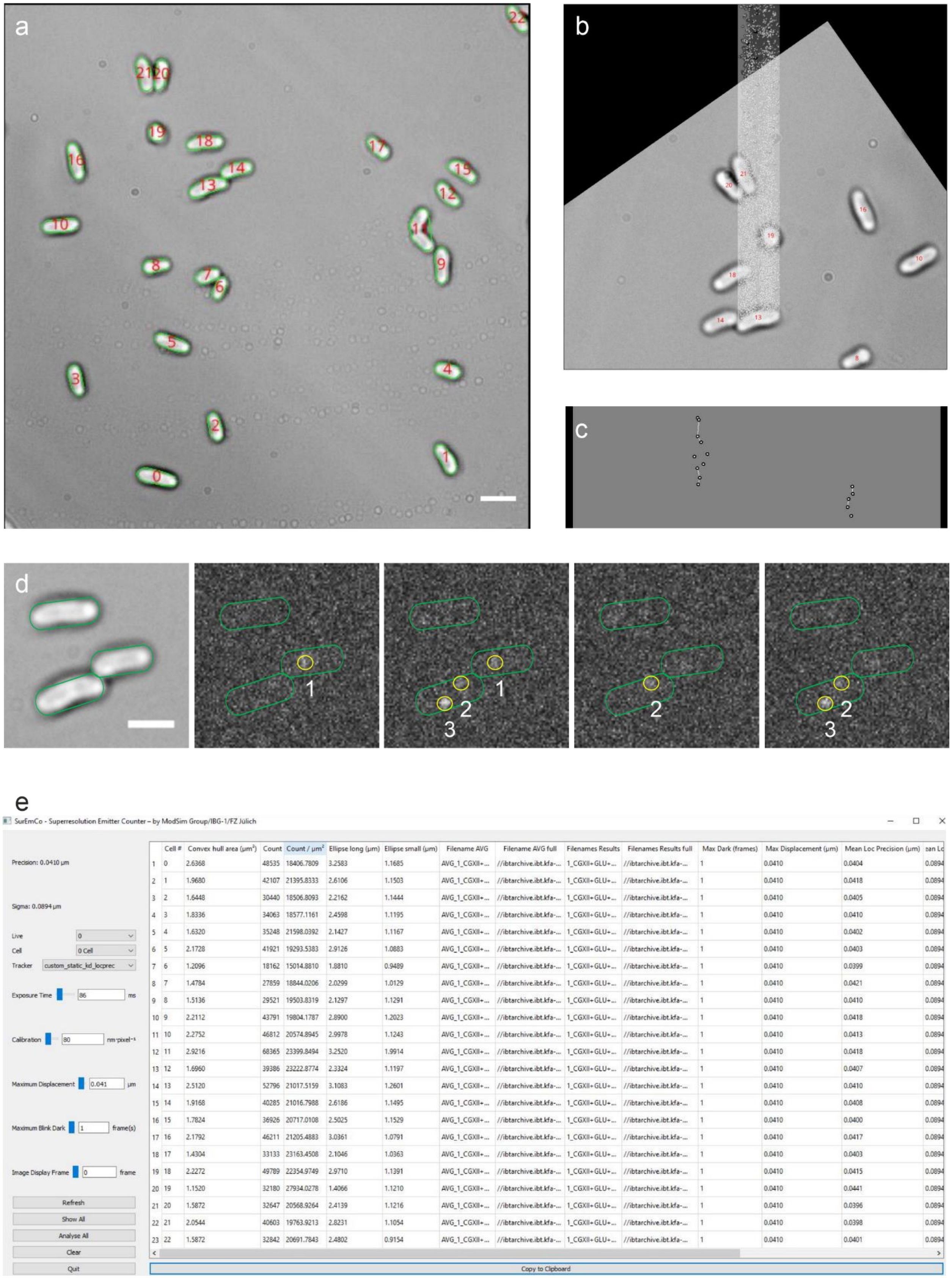


| | Cell # | Convex hull area (µm²) | Count | Count / µm² | Ellipse long (µm) | Ellipse small (µm) | Filename AVG | Filename AVG full | Filenames Results | Filenames Results full | Max Dark (frames) | Max Displacement (µm) | Mean Loc Precision (µm) | ean Lc |
|---|---|---|---|---|---|---|---|---|---|---|---|---|---|---|
| 1 | 0 | 2.6368 | 48535 | 18406.7809 | 3.2583 | 1.1685 | AVG_1_CGXII+... | //ibtarchive.ibt.kfa-... | 1_CGXII+GLU+... | //ibtarchive.ibt.kfa-... | 1 | 0.0410 | 0.0404 | 0.0894 |
| 2 | 1 | 1.9680 | 42107 | 21395.8333 | 2.6106 | 1.1503 | AVG_1_CGXII+... | //ibtarchive.ibt.kfa-... | 1_CGXII+GLU+... | //ibtarchive.ibt.kfa-... | 1 | 0.0410 | 0.0418 | 0.0894 |
| 3 | 2 | 1.6448 | 30440 | 18506.8093 | 2.2162 | 1.1444 | AVG_1_CGXII+... | //ibtarchive.ibt.kfa-... | 1_CGXII+GLU+... | //ibtarchive.ibt.kfa-... | 1 | 0.0410 | 0.0405 | 0.0894 |
| 4 | 3 | 1.8336 | 34063 | 18577.1161 | 2.4598 | 1.1195 | AVG_1_CGXII+... | //ibtarchive.ibt.kfa-... | 1_CGXII+GLU+... | //ibtarchive.ibt.kfa-... | 1 | 0.0410 | 0.0410 | 0.0894 |
| 5 | 4 | 1.6320 | 35248 | 21598.0392 | 2.1427 | 1.1167 | AVG_1_CGXII+... | //ibtarchive.ibt.kfa-... | 1_CGXII+GLU+... | //ibtarchive.ibt.kfa-... | 1 | 0.0410 | 0.0402 | 0.0894 |
| 6 | 5 | 2.1728 | 41921 | 19293.5383 | 2.9126 | 1.0883 | AVG_1_CGXII+... | //ibtarchive.ibt.kfa-... | 1_CGXII+GLU+... | //ibtarchive.ibt.kfa-... | 1 | 0.0410 | 0.0403 | 0.0894 |
| 7 | 6 | 1.2096 | 18162 | 15014.8810 | 1.8810 | 0.9489 | AVG_1_CGXII+... | //ibtarchive.ibt.kfa-... | 1_CGXII+GLU+... | //ibtarchive.ibt.kfa-... | 1 | 0.0410 | 0.0399 | 0.0894 |
| 8 | 7 | 1.4784 | 27859 | 18844.0206 | 2.0299 | 1.0129 | AVG_1_CGXII+... | //ibtarchive.ibt.kfa-... | 1_CGXII+GLU+... | //ibtarchive.ibt.kfa-... | 1 | 0.0410 | 0.0421 | 0.0894 |
| 9 | 8 | 1.5136 | 29521 | 19503.8319 | 2.1297 | 1.1291 | AVG_1_CGXII+... | //ibtarchive.ibt.kfa-... | 1_CGXII+GLU+... | //ibtarchive.ibt.kfa-... | 1 | 0.0410 | 0.0410 | 0.0894 |
| 10 | 9 | 2.2112 | 43791 | 19804.1787 | 2.8900 | 1.2023 | AVG_1_CGXII+... | //ibtarchive.ibt.kfa-... | 1_CGXII+GLU+... | //ibtarchive.ibt.kfa-... | 1 | 0.0410 | 0.0418 | 0.0894 |
| 11 | 10 | 2.2752 | 46812 | 20574.8945 | 2.9978 | 1.1243 | AVG_1_CGXII+... | //ibtarchive.ibt.kfa-... | 1_CGXII+GLU+... | //ibtarchive.ibt.kfa-... | 1 | 0.0410 | 0.0413 | 0.0894 |
| 12 | 11 | 2.9216 | 68365 | 23399.8494 | 3.2520 | 1.9914 | AVG_1_CGXII+... | //ibtarchive.ibt.kfa-... | 1_CGXII+GLU+... | //ibtarchive.ibt.kfa-... | 1 | 0.0410 | 0.0418 | 0.0894 |
| 13 | 12 | 1.6960 | 39386 | 23222.8774 | 2.3324 | 1.1197 | AVG_1_CGXII+... | //ibtarchive.ibt.kfa-... | 1_CGXII+GLU+... | //ibtarchive.ibt.kfa-... | 1 | 0.0410 | 0.0407 | 0.0894 |
| 14 | 13 | 2.5120 | 52796 | 21017.5159 | 3.1083 | 1.2601 | AVG_1_CGXII+... | //ibtarchive.ibt.kfa-... | 1_CGXII+GLU+... | //ibtarchive.ibt.kfa-... | 1 | 0.0410 | 0.0410 | 0.0894 |
| 15 | 14 | 1.9168 | 40285 | 21016.7988 | 2.6186 | 1.1495 | AVG_1_CGXII+... | //ibtarchive.ibt.kfa-... | 1_CGXII+GLU+... | //ibtarchive.ibt.kfa-... | 1 | 0.0410 | 0.0408 | 0.0894 |
| 16 | 15 | 1.7824 | 36926 | 20717.0108 | 2.5025 | 1.1529 | AVG_1_CGXII+... | //ibtarchive.ibt.kfa-... | 1_CGXII+GLU+... | //ibtarchive.ibt.kfa-... | 1 | 0.0410 | 0.0400 | 0.0894 |
| 17 | 16 | 2.1792 | 46211 | 21205.4883 | 3.0361 | 1.0791 | AVG_1_CGXII+... | //ibtarchive.ibt.kfa-... | 1_CGXII+GLU+... | //ibtarchive.ibt.kfa-... | 1 | 0.0410 | 0.0417 | 0.0894 |
| 18 | 17 | 1.4304 | 33133 | 23163.4508 | 2.1046 | 1.0363 | AVG_1_CGXII+... | //ibtarchive.ibt.kfa-... | 1_CGXII+GLU+... | //ibtarchive.ibt.kfa-... | 1 | 0.0410 | 0.0403 | 0.0894 |
| 19 | 18 | 2.2272 | 49789 | 22354.9749 | 2.9710 | 1.1391 | AVG_1_CGXII+... | //ibtarchive.ibt.kfa-... | 1_CGXII+GLU+... | //ibtarchive.ibt.kfa-... | 1 | 0.0410 | 0.0415 | 0.0894 |
| 20 | 19 | 1.1520 | 32180 | 27934.0278 | 1.4066 | 1.1210 | AVG_1_CGXII+... | //ibtarchive.ibt.kfa-... | 1_CGXII+GLU+... | //ibtarchive.ibt.kfa-... | 1 | 0.0410 | 0.0441 | 0.0894 |
| 21 | 20 | 1.5872 | 32647 | 20568.9264 | 2.4139 | 1.1216 | AVG_1_CGXII+... | //ibtarchive.ibt.kfa-... | 1_CGXII+GLU+... | //ibtarchive.ibt.kfa-... | 1 | 0.0410 | 0.0396 | 0.0894 |
| 22 | 21 | 2.0544 | 40603 | 19763.9213 | 2.8231 | 1.1054 | AVG_1_CGXII+... | //ibtarchive.ibt.kfa-... | 1_CGXII+GLU+... | //ibtarchive.ibt.kfa-... | 1 | 0.0410 | 0.0398 | 0.0894 |
| 23 | 22 | 1.5872 | 32842 | 20691.7843 | 2.4802 | 0.9154 | AVG_1_CGXII+... | //ibtarchive.ibt.kfa-... | 1_CGXII+GLU+... | //ibtarchive.ibt.kfa-... | 1 | 0.0410 | 0.0401 | 0.0894 |

**Figure S2. Ribosome counting with the SurEmCo software.** *C. glutamicum* cells were cultivated in CGXII+GLU+CAA until mid-exponential phase, prepared for SMLM and imaged as described in materials and methods. **a** – Cell regions are identified by the software. Each identified cell region is depicted on the original transmission image (green line delimitates the cell perimeter) and attributed an individual numerical identifier (red number). Scale bar – 3 µm. **b** – Emitters identified by the software within the cell boundaries are tracked in consecutive frames (3D projection). **c** – Zoom in on the 3D projection. Emitters (dots) that appear at the same position within a set precision threshold in subsequent frames are considered the same emitter (shown connected by a white line), and therefore counted as a single emitter. **d** – Examples of emitters counted as single events. #1 and #2 appear at the same position in consecutive frames. Likewise, since we chose a maximum blink dark value of one frame for all our analyses, emitter #3 was also counted as a single molecule. Scale bar – 2 µm. **e** – Analysis parameters (left column) can be adjusted and results are displayed per cell on the right. SurEmCo is a Python tool that is available under the BSD license at https://github.com/modsim/suremco.

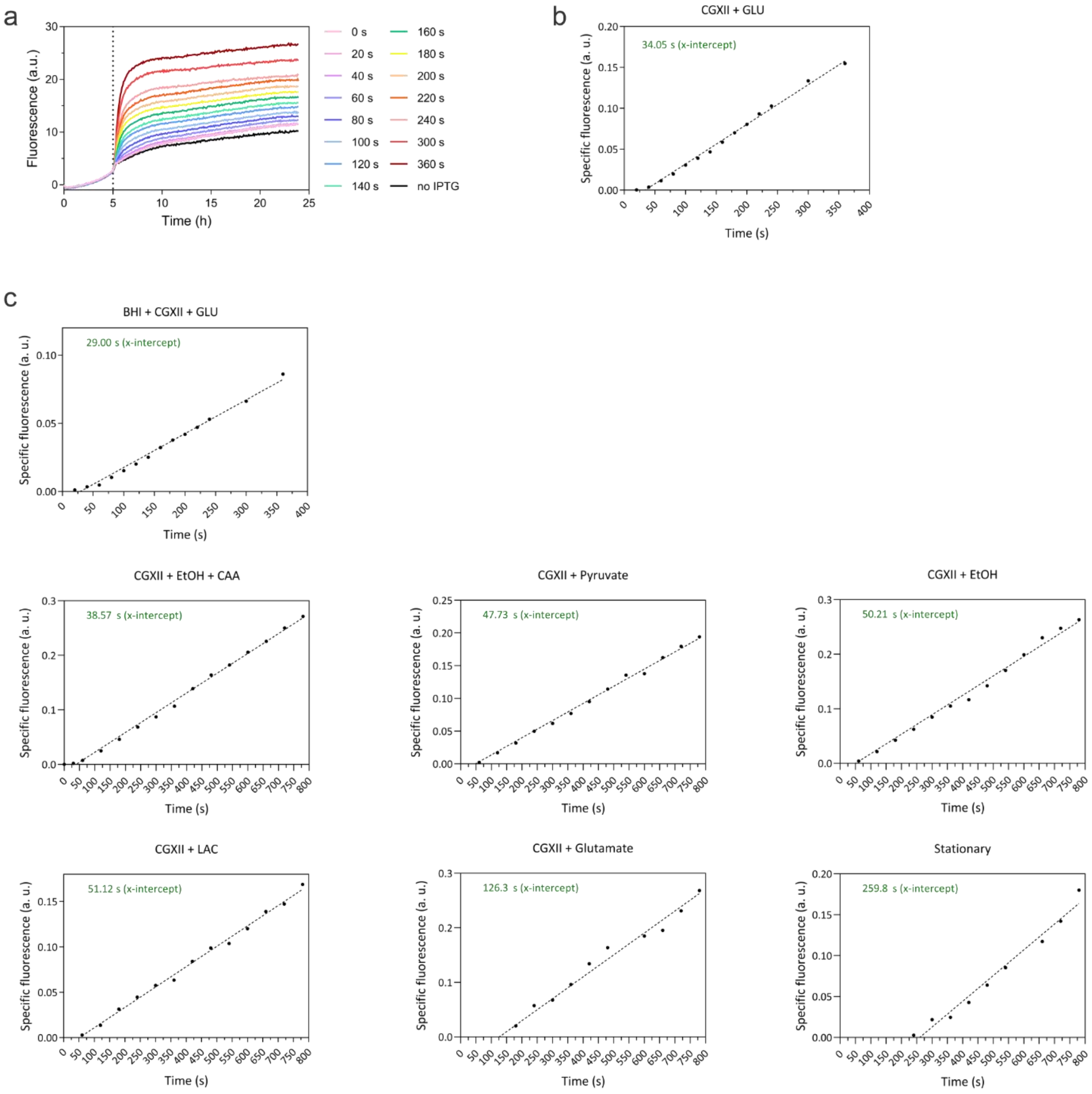


**Figure S3. Fluorescent assay for determination of the translation elongation rate. a** – Strain *C. glutamicum* MB001(DE3)-pMKEx2-*eyfp* was grown in CGXII+GLU. At mid-exponential phase *eyfp* expression was induced with IPTG (vertical dashed line). Translation was stopped by chloramphenicol addition at 20-60 s intervals in a robotics platform. **b** - The specific fluorescence of a 3 h interval after the fluorescent signal reached a constant maximum was plotted for each time point after normalization with the no IPTG control. A linear regression was used to determine the intersection point on the *x*-axis. An EYFP (238 aa) fluorescence signal above the control is detected after ~34 s. Assuming the initial translation initiation steps are similar to *E. coli* and last as long (~10 s), 3 s were deducted from the x-intercept values (the extra 7 s are inherent to our robotic setup), resulting in a translation elongation rate of ~7.66 aa $s^{-1}$ in this case. **c** - Examples of translation elongation assays for all conditions tested.

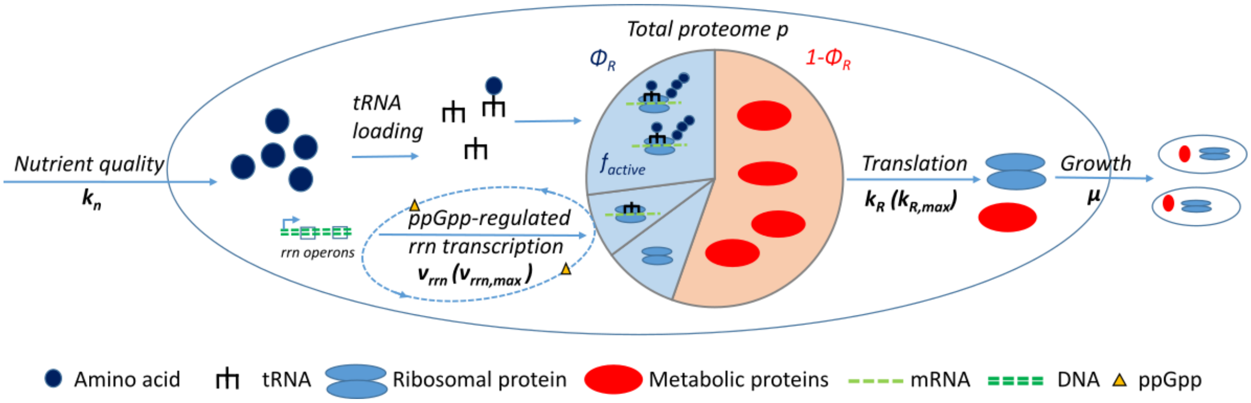


**Figure S4. Schematic of the coarse-grained self-replicator model and its most important functions.** In brief, amino acids are available in the cell as precursors for protein translation ($k_R$) depending on the nutrient quality ($k_n$), i.e. the nutrient condition of the cell. Complexes of loaded tRNAs and ribosomes constitute the fraction of actively translating ribosomes ($f_{active}$) within the fraction of ribosomal proteins ($\Phi_R$) in the cell. Translated proteins are diluted into newly formed biomass expressed by the specific growth rate $\mu$. The governing principle behind the self-replicator model is rooted in the finite resource effect: too few ribosomes limit growth while too many of them increase the investment in amino acids that are, in turn, not available for growth either. This reflects the cells' "balancing act" between amino acid flux and protein translation capacity and makes the self-replicator model well suited to describe the phenomenological Rb/μ correlation. In the model, a ppGpp regulatory control mechanism senses suboptimal growth states. The control mechanism initiates compensatory *rrn* gene regulation to fine-tune ribosome production ($v_{rrn}$) and restore optimal growth. Calibrated with experimental data of *C. glutamicum* at hand, the model delivers quantitative insights into cellular key parameters that are hitherto inaccessible or difficult to obtain experimentally ($k_R$, $v_{rnn}$).

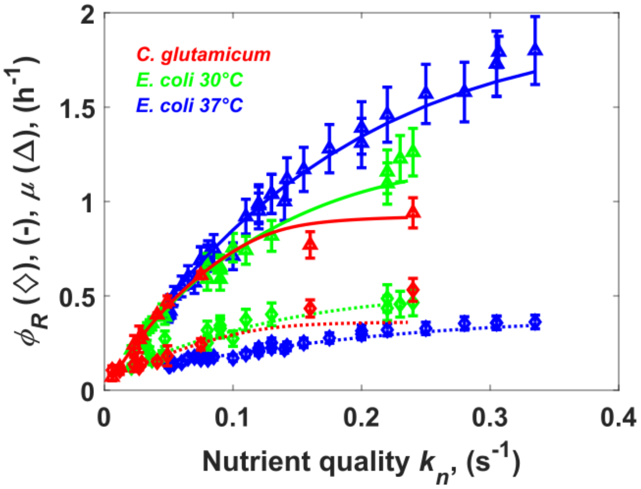


**Figure S5. Result of model calibration for *C. glutamicum* and *E. coli*.** Model fits of specific growth rates $\mu$ (solid lines, 0.06 -1.8 $h^{-1}$) and ribosomal fractions $\Phi_R$ (dotted lines) over the range of $k_n$ values for *C. glutamicum* (red, 0.006-0.240 $s^{-1}$), *E. coli* 30 °C (green, 0.021-0.240 $s^{-1}$) and *E. coli* 37 °C (blue, 0.050-0.335 $s^{-1}$). Experimental data used for calibrating the model are reported in Table S9 for *C. glutamicum*. For *E. coli*, data at 30 °C see [17, 24] and for 37 °C see [10, 14, 15, 18]. Data used for model calibration is also contained in the Supplementary Model File. For the model calibration procedure see Material and Methods and Supplementary Note 5C.

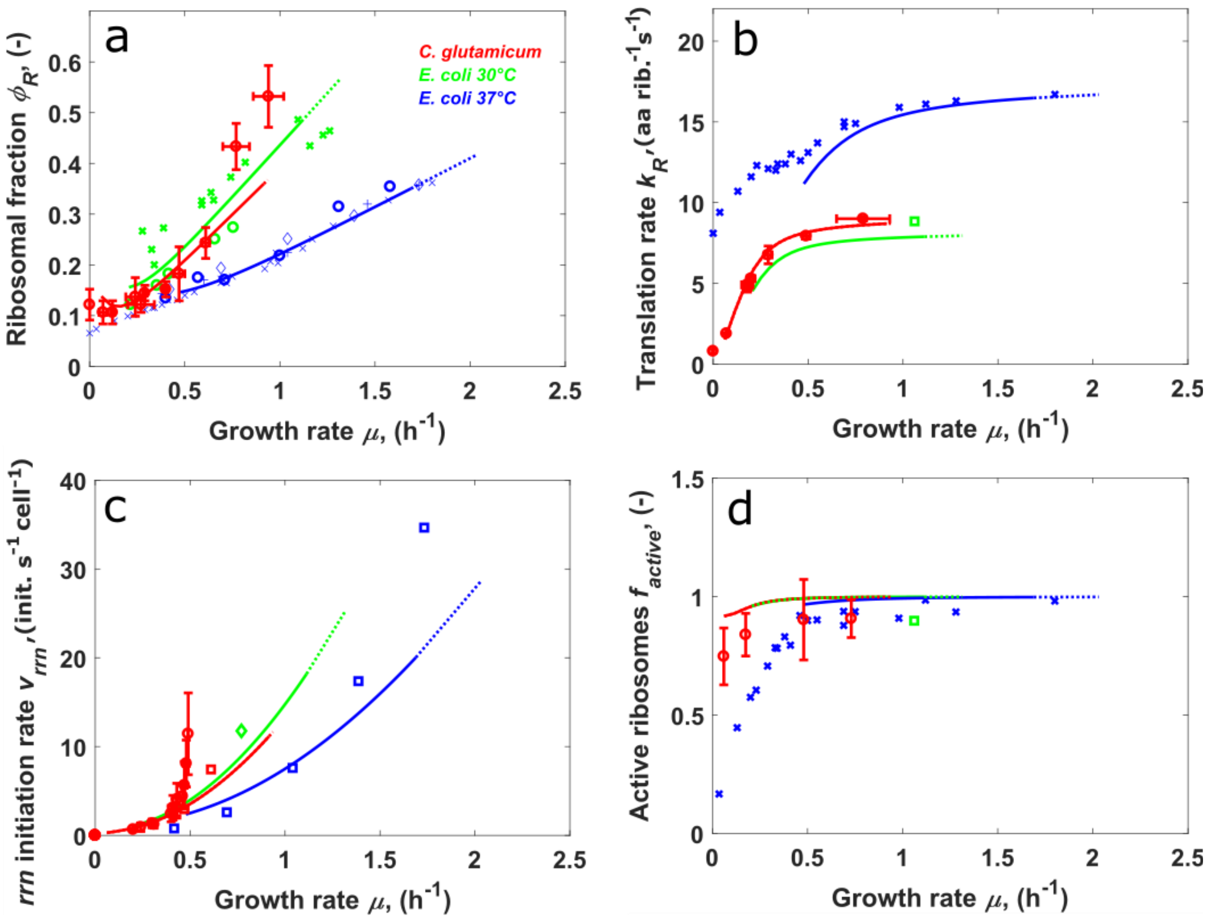


**Figure S6. Comparison of the Rb/µ correlation, inferred rates and active ribosome fraction observed in *C. glutamicum* and *E. coli.*** **a -** Phenomenological observation of the Rb/µ correlation for *C. glutamicum* (this work), *E. coli* grown at 30 °C and *E. coli* grown at 37 °C. Courses of ribosomal protein fractions $\Phi_R$ are shown at various growth rates. Experimental data is shown as symbols (see Table S2 for *C. glutamicum* and Supplementary Model File for *E. coli*). *E. coli* experimental data was taken from the literature: (**o**) [15], (**+**) [18], (**◊**) [10], (**x**) [14], (**o**) [24], (**x**) [17]. Solid lines represent simulations of the respective simulated values using the calibrated model with estimated parameters (see Table S6): *C. glutamicum* (red), *E. coli* 30 °C (green) and *E. coli* 37 °C (blue), respectively. **b** - Comparison of experimental and of model-derived translation rates $k_R$. *E. coli* experimental data was retrieved from the literature: (□) [19], (**x**) [14]. Translation elongation rates of *C. glutamicum* are listed in Table S3. **c -** Comparison of experimental and model-derived ribosome production rates (i.e. *rrn* operon initiation rates). *C. glutamicum* data was calculated from SMLM data (□) (see Supplementary Note 2) and from R/P data (**o**) (Table S5). *E. coli* experimental data was calculated from (□) [10] and (**◊**) [20]. The rate of *rrn* operon transcription initiation $v_{rrn}$ was calculated according to $v_{rrn} = N_R \cdot \mu$ [10] and was experimentally determined by measuring the number of ribosomes per cell $N_R$ at the corresponding specific growth rate $\mu$ ($s^{-1}$).

$v_{rrn}$ is also termed ribosome production rate because it is assumed that each *rrn* transcription initiation triggers the production of one extended ribosome. **d -** Comparison of experimental and model-derived fractions $f_{active}$ of actively translating ribosomes. For *E. coli* 37 °C experimental raw data (corresponding sets of $\mu$, $k_R$, total RNA per total protein (w/w)) were taken from [14] and fractions of active ribosomes (**x**) calculated according to Supplementary Note 4 assuming $\sigma'_{E.\ coli(37°C)} = m_{RNA} /(m_{aa} \cdot 0.86) = 1.56*10^4 \approx \sigma'_{E.\ coli(30°C)}$ [14]. For *E. coli* 30 °C, experimental raw data for $k_R$ and $\mu$ were taken from [19], and the fraction of total RNA per total protein was recalculated from $\Phi_{R,sim}$ at $\mu = 1.06\ h^{-1}$ (see **a**) for calculation of the active ribosome fraction (□). Dashed lines represent growth under "super-rich medium" represented by $k_n$ increased by two orders of magnitude (Table S7).

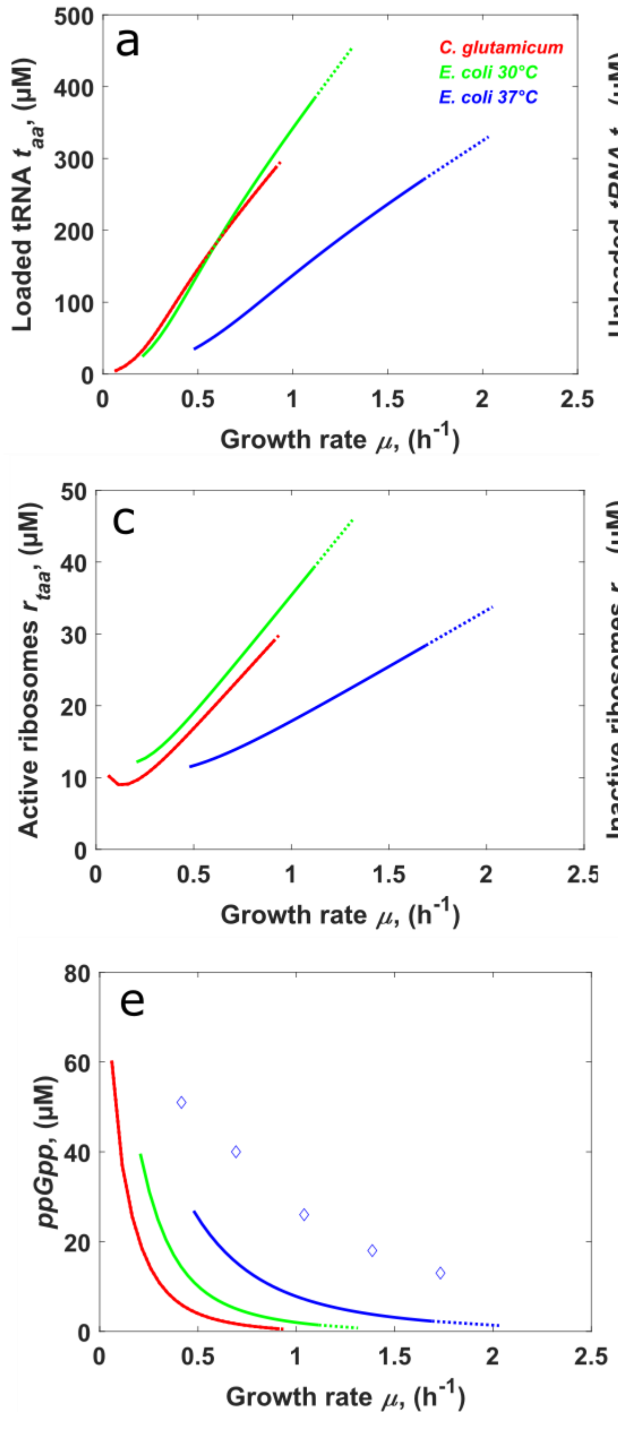


**Figure S7. Model-inferred tRNA loading status, ribosome activity and ppGpp concentrations across growth rates for *C. glutamicum* and *E. coli*.** This figure summarizes, across growth rates, concentrations of **a –** aminoacylated tRNA ($t_{aa}$), **b –** unloaded tRNA ($t_f$), **c –** complexes of actively translating ribosomes ($r_{taa}$), **d –** non-translating complexes due to bound unloaded tRNA ($r_{tf}$), **e –** *ppGpp*. Experimental data (◊) is taken from [10]. Dashed lines represent growth under “super-rich” medium represented by $k_n$ increased by two orders of magnitude (see Table S7).

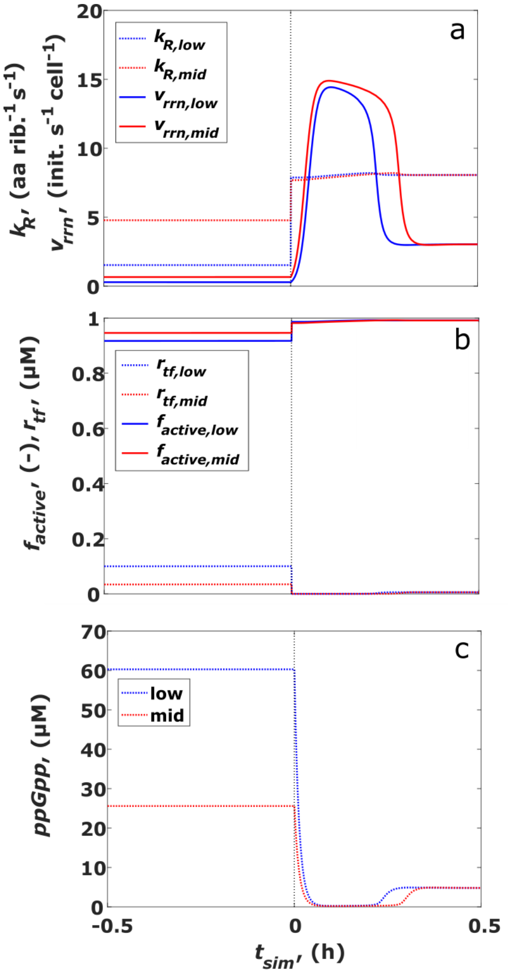


**Figure S8. Simulation of the upshift experiment – interrogation of the model.** Comparison of inferred translation elongation as well as ribosome production rates **(a),** instantaneous concentrations of inactive (unloaded tRNA-) ribosome complexes and active ribosome fractions $f_{active}$ **(b)** and the ppGpp concentration (**c**). The nutrient shift (vertical dashed line at $t$=0) was initiated by an increase of the nutrient quality parameter (see Materials and Methods) either from $k_{n,glutamate}$ = 0.006 s$^{-1}$ (index "low", blue curves) or $k_{n,EtOH}$ = 0.016 s$^{-1}$ (index "mid", red curves) to $k_{n,glucose}$ = 0.051 s$^{-1}$ (Table S9). The corresponding instantaneous growth rates and ribosome fractions are found in Fig. 5d.
